# Rational control of structural off-state heterogeneity in a photoswitchable fluorescent protein provides switching contrast enhancement

**DOI:** 10.1101/2021.11.05.462999

**Authors:** Virgile Adam, Kyprianos Hadjidemetriou, Nickels Jensen, Robert L. Shoeman, Joyce Woodhouse, Andrew Aquila, Anne-Sophie Banneville, Thomas R. M. Barends, Victor Bezchastnov, Sébastien Boutet, Martin Byrdin, Marco Cammarata, Sergio Carbajo, Nina Eleni Christou, Nicolas Coquelle, Eugenio De la Mora, Mariam El Khatib, Tadeo Moreno Chicano, R. Bruce Doak, Franck Fieschi, Lutz Foucar, Oleksandr Glushonkov, Alexander Gorel, Marie Luise Grünbein, Mario Hilpert, Mark Hunter, Marco Kloos, Jason E. Koglin, Thomas J. Lane, Mengning Liang, Angela Mantovanelli, Karol Nass, Gabriela Nass Kovacs, Shigeki Owada, Christopher M. Roome, Giorgio Schirò, Matthew Seaberg, Miriam Stricker, Michel Thépaut, Kensuke Tono, Kiyoshi Ueda, Lucas M. Uriarte, Daehyun You, Ninon Zala, Tatiana Domratcheva, Stefan Jakobs, Michel Sliwa, Ilme Schlichting, Jacques-Philippe Colletier, Dominique Bourgeois, Martin Weik

## Abstract

Reversibly photoswitchable fluorescent proteins are essential markers for advanced biological imaging, and optimization of their photophysical properties underlies improved performance and novel applications. Here we establish a link between photoswitching contrast, a key parameter that largely dictates the achievable resolution in nanoscopy applications, and chromophore conformation in the non-fluorescent state of rsEGFP2, a widely employed label in REversible Saturable OpticaL Fluorescence Transitions (RESOLFT) microscopy. Upon illumination, the *cis* chromophore of rsEGFP2 isomerizes to two distinct *off*-state conformations, *trans1* and *trans2*, located on either side of the V151 side chain. Reducing or enlarging the side chain at this position (V151A and V151L variants) leads to single *off*-state conformations that exhibit higher and lower switching contrast, respectively, compared to the rsEGFP2 parent. The combination of structural information obtained by serial femtosecond crystallography with high-level quantum chemical calculations and with spectroscopic and photophysical data determined *in vitro* suggests that the changes in switching contrast arise from blue- and red-shifts of the absorption bands associated to *trans1* and *trans2*, respectively. Thus, due to elimination of *trans2*, the V151A variants of rsEGFP2 and its superfolding variant rsFolder2 display a more than two-fold higher switching contrast than their respective parent proteins, both *in vitro* and in *E. coli* cells. The application of the rsFolder2-V151A variant is demonstrated in RESOLFT nanoscopy. Our study rationalizes the connection between structural and photophysical chromophore properties and suggests a means to rationally improve fluorescent proteins for nanoscopy applications.

## Introduction

Reversibly switchable fluorescent proteins (RSFPs ^[1]^) are photochromic markers that are key to multiple super-resolution microscopy (nanoscopy) schemes including RESOLFT (Reversible Saturable Optical Fluorescence Transition ^[2]^), NL-SIM (Non Linear Structured Illumination Microscopy ^[3]^), SOFI (Super resolution Optical Fluctuation Imaging ^[4]^) and multicolor PALM (Photo Activated Localization Microscopy ^[5]^). They are also central tools in contrast enhancing approaches such as OLID (Optical Lock-In Detection microscopy ^[6]^) and in single channel multicolor approaches such as OPIOM (Out-of-Phase Imaging after Optical Modulation ^[7]^). With the exception of Dreiklang ^[2c]^ and its variant SPOON ^[8]^, RSFPs typically switch between a fluorescent *on*- and a non-fluorescent *off*-state through light-induced *cis*-*trans* isomerization of their p-hydroxybenzylidene imidazolinone chromophore ^[1a, 9]^. Depending on whether the same wavelength that induces fluorescence switches the RSFP from the *on*-to the *off*-state or *vice versa*, they are said to be negative or positive, respectively ^[10]^. At neutral pH, the chromophore of negative RSFPs (such as rsEGFP2 and its rsFolder variants studied here) is generally *cis*-anionic in the *on*-state and *trans*-protonated in the *off*-state, although exceptions have been reported in rsGamillus ^[11]^. *Off*-switching is promoted by illumination with wavelengths near the fluorescence excitation maximum (typically 488 nm) while *on*-switching requires illumination in the near UV region (typically 405 nm, Fig. 1a).

**Figure 1:**
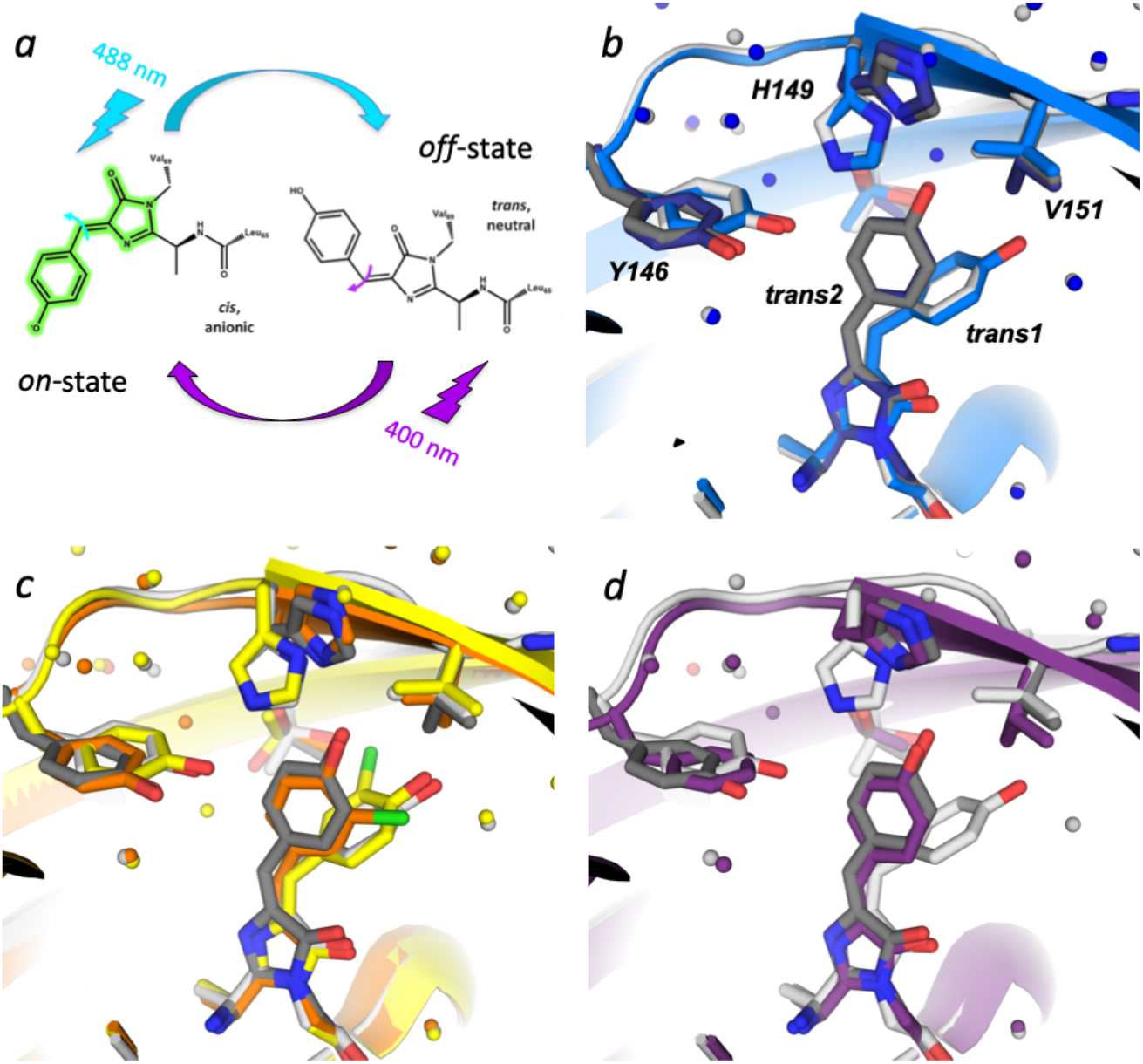
Photoswitching and *off*-state conformations in parental rsEGFP2. (*a*) rsEGFP2 can be photoswitched from the fluorescent *on*-state (anionic *cis* chromophore) to the non-fluorescent *off*-state (neutral trans chromophore) by illumination with 488 nm light, and back by illumination with 405-nm light. Photoswitching involves chromophore isomerization (blue and purple arrows on chromophore methylene bridge) and a change in protonation state of the phenol group. (*b*) Structures of parental rsEGFP2 in its *off*-state solved from RT SFX data. *Off*-state models of parental rsEGFP2 solved from RT SFX data earlier (PDB entry 6T39^[18]^) and in this work (PDB entry 7O7U). *Trans1* and *trans2* are occupied at 40% (light grey) and 30% (dark grey) in parental rsEGFP2 (this work) and at 65% (light blue) and 25% (dark blue) in 6T39, respectively. (*c, d*) *Off*-state models of parental rsEGFP2 solved from RT SFX data in this work (light and dark grey), overlaid with *trans* conformations in synchrotron structures of rsEGFP2 containing a monochlorinated chromophore ^[19]^ solved from crystals with looser (yellow) and tighter (orange) crystal packings ^[19]^ (*c*) and with the o*ff*-state model of rsFolder (PDB entry 5DU0^[21]^) in purple (*d*).

In all nanoscopy applications relying on RSFPs as labels, image quality and the achievable spatial resolution are mainly determined by the following photophysical characteristics ^[10]^: i) the fluorescence brightness, being defined as the product of the extinction coefficient of the *on*-state and the fluorescence quantum yield, ii) the ensemble switching speed, *i*.*e*. the time required to switch the ensemble from the *on*-to the *off*-state, or *vice versa*, iii) the switching fatigue, being defined as the fraction of an RSFP ensemble being photobleached per full switching cycle, and iv) the switching contrast, that is, the ratio between the fluorescence signal after *on*-switching and the residual signal after *off*-switching. Such residual fluorescence after *off*-switching mainly originates from back switching of the *off*-state chromophore by the *off*-switching light ^[12]^. Among these characteristics, a high switching contrast is most critical for achieving high spatial resolution, and engineering efforts have thus been recently conducted to maximize it ^[11, 13]^. If we neglect the possibility that the chromophore is not fully in the fluorescent *cis*-anionic state after *on*-switching ^[14]^, the switching contrast is a function of the *on*-to-*off* and *off*-to-*on* switching quantum yields and the extinction coefficients of both the *on*- and *off*-states at the *off*-switching wavelength (see eq. 1 in the Results section). In recent experimental ^[15]^ and computational ^[16]^ studies, it was proposed that the switching contrast is controlled by the relative stability of RSFPs in their *on-* and *off*-states *via* the number of hydrogen bonds between the chromophore and the protein pocket and its water molecules in each state. Here, focusing on rsEGFP2 and its rsFolder variants, we expand this view by showing how different conformations of the *off*-state chromophore modulate switching contrast, opening the door to rational optimization of RSFPs for enhanced nanoscopy applications.

rsEGFP2 (Figure 1a) has been generated based on EGFP and is widely applied in RESOLFT microscopy thanks to its fast maturation and favorable balance between fluorescence brightness, switching quantum yields, photofatigue resistance and switching contrast ^[17]^. Recently, evidence for conformational heterogeneity in the *off*-state of rsEGFP2 at room temperature was provided by serial femtosecond crystallography (SFX), where in addition to the major *trans* state (*trans1* conformer), a hitherto unobserved *trans* isomer (*trans2* conformer) was observed ^[18]^, which displays different twist and tilt dihedral angles (φ and τ dihedral angles; see also Supplementary table S3), protein environment and H-bonding network (Supplementary figure S1). Interestingly, cryo-crystallographic synchrotron data revealed a similar *trans2* isomer upon *off*-switching of an rsEGFP2 variant containing a monochlorinated chromophore when crystals with a contracted unit cell were examined, whereas a conformation similar to *trans1* was populated in crystals with a larger unit cell ^[19]^. *Trans1* and *trans2* conformers are located on either side of the V151 side chain (Supplementary figure S1), a residue that needs to transiently retract for the chromophore to switch between its *cis* and *trans* conformations (see supplementary movie in ^[20]^) as suggested by time-resolved SFX ^[20]^. To start addressing the role of this residue in photoswitching, a variant with more space was generated by mutating V151 to an alanine ^[20]^. During the preliminary photophysical characterization ^[20]^ neither the switching contrast or the structure of the rsEGFP2-V151A variant was analyzed nor was the sEGFP2-V151L control with a more bulkier side chain generated.

The *trans2* conformation in rsEGFP2 is also very similar to the *trans* conformation observed in the *off*-state of rsFolder, a superfolding variant of rsEGFP2 that harbors a phenylalanine instead of a tyrosine at position 146 and that was designed to facilitate RESOLFT microscopy in “hostile” environments such as the bacterial periplasm ^[21]^. Strikingly, rsFolder has a much lower switching contrast than rsEGFP2 and rsFolder2, a single mutant of rsFolder with a tyrosine at position 146 and thus the same chromophore pocket as rsEGFP2 ^[21]^. This observation suggests a possible link between the presence of *trans2* and a reduced switching contrast, raising the intriguing hypothesis that removal of the *trans2* fraction in rsEGFP2 could further enhance its switching contrast.

Here, we investigate the structural *off*-state heterogeneity in rsEGFP2 and eliminated it in V151A and V151L variants by shortening or lengthening the amino-acid side chain at position 151, respectively. The V151A variant exhibits only the *trans1* conformation and a substantially higher switching contrast compared to the rsEGFP2 parent, whereas the V151L variant shows only the *trans2* conformation and a lower contrast. The effects of the V151A and V151L mutations on the switching contrast are reproduced in rsFolder2. We show that changes in switching contrast between the investigated variants mainly result from differences in extinction coefficients of the corresponding *off*-states at the *off*-switching illumination wavelength. Furthermore, we postulate rapid exchange dynamics between *trans1* and *trans2* in parental rsEGFP2 and in rsFolder2 and suggest that the achieved equilibrium depends on environmental factors, a notion we refer to as “photoswichting fragility”. Finally, we show that the V151A variants maintain their gain in switching contrast *in vivo* and investigate the potential of rsFolder2-V151A for RESOLFT nanoscopy.

## Results

### Structural heterogeneity in the off-state of parental rsEGFP2

In order to corroborate the observation of a second *trans* isomer in parental rsEGFP2 ^[18]^, a follow-up SFX experiment was carried out at the Linac Coherent Light Source (LCLS) using microcrystals of parental rsEGFP2 of the same crystal batch used earlier ^[18, 20]^. The *on*-state crystals were photoswitched by 488 nm light ^[22]^ and the room-temperature (RT) structure of the resulting *off*-state solved at 1.7 Å resolution (PDB entry 7O7U; see SI for details). Both *trans1* and *trans2* chromophore conformations are again present (Figure 1b, Supplementary figure S2). They agree well with those observed earlier ^[18]^, as well as with those observed in cryo-crystallographic structures of an rsEGFP2 variant containing a monochlorinated chromophore ^[19]^ (Figure 1c, Supplementary table S3). The *trans2* conformation is similar to the one adopted by rsFolder in its *off*-state ^[21]^ (Figure 1d).

### Structural heterogeneity bisected in the off-states of rsEGFP2 V151A and V151L variants

Given that the *trans1* and *trans2* chromophore conformations lie on either side of the V151 side chain (Figure 1b-d), we reasoned that this residue could also control the *off*-state heterogeneity. In addition to the rsEGFP2 variant with a shortened side chain (V151A ^[20]^), one with an enlarged (V151L) side chain was therefore generated and non-fluorescent *off*-state structures of both solved from RT SFX data collected at the SPring-8 Angstrom Compact free electron Laser (SACLA) from microcrystals after 488 nm light illumination ^[22]^ (Figure 2, Supplementary figure S3, Supplementary table S2). The *off*-state structures (V151A: PDB entry 7O7X, V151L: PDB entry 7O7W) display only one chromophore conformation: *trans1* for V151A (Figure 2a) and *trans2* for V151L (Figure 2b). Absorption spectroscopy indicates that about 85% and 77% of microcrystalline rsEGFP2-V151A and -V151L chromophores have switched from the *on*-to the *off*-state, respectively (Supplementary figure S4). *Trans1* and *trans2* chromophore conformations were modelled at 80% (75%) occupancy in rsEGFP2-V151A (V151L) and the residual *cis* conformer at 20% (25%). Spectroscopic and crystallographic *on*- and *off*-state occupancies are thus consistent. In addition to differences in chromophore conformations, the *off*-states of the two variants also differ in their His149 and Tyr146 conformations. In the V151A variant His149 forms a hydrogen bond with Tyr146 and the *trans1* chromophore forms a hydrogen bond with a water molecule (distance: 2.7 Å, Supplementary figure S1c, Figure 2a), whereas in the V151L variant His149 is hydrogen bonded to the *trans2* chromophore (distance of 2.5 Å between the chromophore phenol group and His149ND1; Supplementary figure S1b, Figure 2b). Synchrotron cryo-crystallography structures of rsEGFP2-V151A (*off*-state: PDB entry 7O7C, *on*-state: PDB entry 7O7D) and - V151L (*off*-state: PDB entry 7O7H, *on*-state: PDB entry 7O7E) also feature a *trans1* and *trans2* chromophore in their *off*-state, respectively (Supplementary figure S5b, d) and a *cis* chromophore in the *on*-state (Supplementary figure S5a, c).

**Figure 2:**
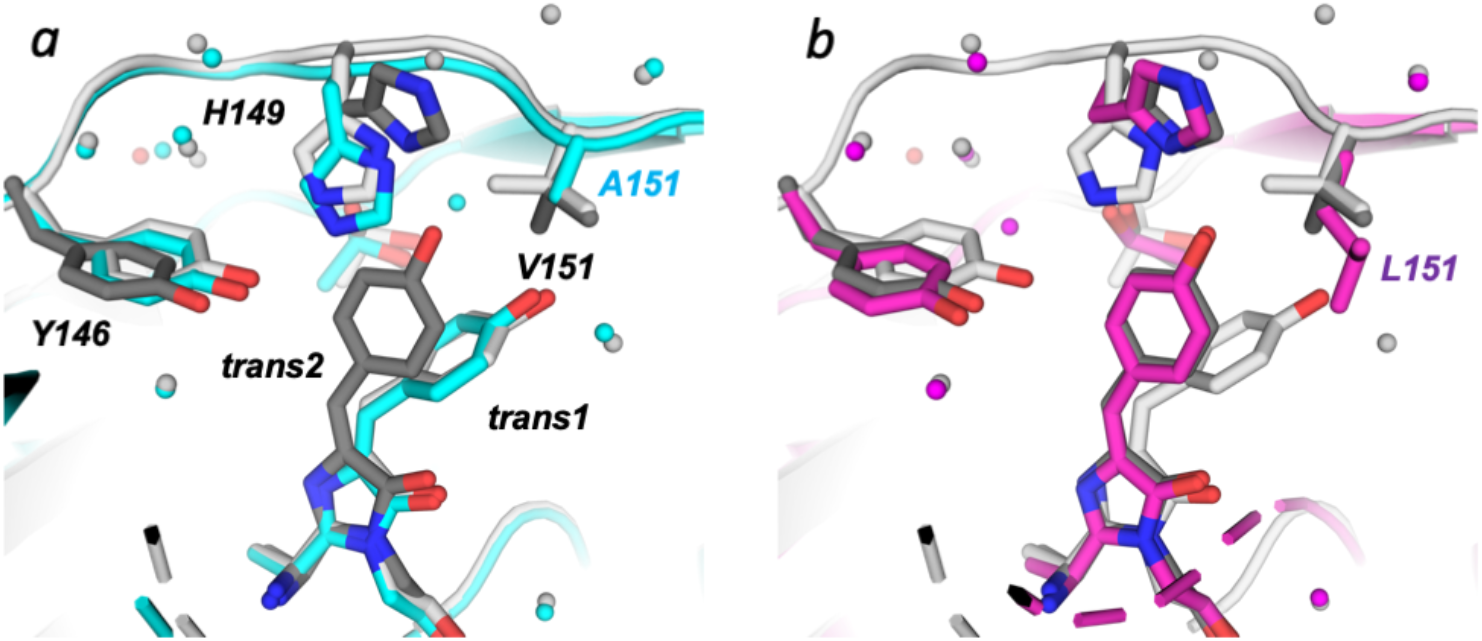
Structures of rsEGFP2 and its V151A and V151L variants in their *off*-states solved from RT SFX data. *Off*-state models of (a) rsEGFP2-V151A (cyan; PDB entry 7O7X) and (b) -V151L (purple; PDB entry 7O7W) variants are superimposed on the model of parental rsEGFP2 in the *off*-state solved from RT SFX data (PDB entry 7O7U), featuring *trans1* in light grey and *trans2* in dark grey. *Trans1* and *trans2* are occupied at 80% and 75% in rsEGFP2 V151A and V151L, respectively. The *cis* conformers were removed for clarity.

Overall, the RT SFX structures strongly suggest that the conformational *off*-state heterogeneity (*trans1, trans2*) seen in parental rsEGFP2 is eliminated in the rsEGFP2-V151A and -V151L variants, with *trans1* being occupied in the former and *trans2* in the latter. Thus, the residue at position 151 controls the *off*-state chromophore conformations (for a discussion of the modulation of conformational off-state heterogeneity see Supplementary Text S4).

### Occupancies of trans1 and trans2 conformations in parental rsEGFP2 and rsFolder2 are sensitive to experimental conditions

A puzzling observation is that the relative occupancies of the *trans1* and *trans2* conformations differ in *off-*state crystal structures of rsEGFP2 determined from three different SFX data sets, although the same batch of microcrystals was used (Supplementary text S1). In addition, attempts to observe the *trans2* conformation in macrocrystals of parental rsEGFP2 upon RT illumination at various intensities by synchrotron cryo-crystallography remained unsuccessful (see Supplementary text S2). In contrast, the rsFolder2 *off*-state structure determined by synchrotron cryo-crystallography (PDB entry 7AMF) showed residual occupancy of the *trans2* chromophore in addition to a mainly occupied *trans1* (Supplementary figure S6), similarly to the parental rsEGFP2 structures determined by RT SFX. These results suggest that relative occupancies of the two *off*-state conformations may change depending on even subtle differences in experimental conditions (see also ^[19]^). In contrast, the rsEGFP2-V151A and -V151L structures in their *off-*states determined by SFX (Figure 2) are similar to those derived from cryo-crystallographic data we collected from macrocrystals at the European Synchrotron Radiation Facility (ESRF) (Supplementary table S1; Supplementary figure S5).

### Determination of switching contrast, switching quantum yields and extinction coefficients of rsEGFP2, rsFolder2 and their variants embedded in polyacrylamide gels

To explore a possible correlation between the *off*-state chromophore conformations and the switching contrast, we measured the switching kinetics of rsEGFP2-V151A, rsEGFP2-V151L and parental rsEGFP2 that contain either *trans1, trans2*, or both conformations, respectively. We also investigated rsFolder and rsFolder2, as well as the two variants rsFolder2-V151A and rsFolder2-V151L.

We embedded the seven investigated variants in polyacrylamide gels and recorded their fluorescence switching curves under laser illumination at 488 nm, using a wide field fluorescence microscope (Figure 3). The switching contrast was calculated as the ratio of the initial fluorescence in the *on*-state after illumination at 405 nm divided by the residual steady-state fluorescence after *off*-switching with 488 nm light (Table 1). We found that for both rsEGFP2 and rsFolder2 the switching contrast is reduced (× ∼0.33) in the V151L variants and increased (× ∼2.6) in the V151A variants compared to the parent proteins. The switching contrasts measured in the V151L variants (∼15) are similar to that of rsFolder (∼20), whereas that measured for both V151A variants exceeds 100.

**Table 1.**
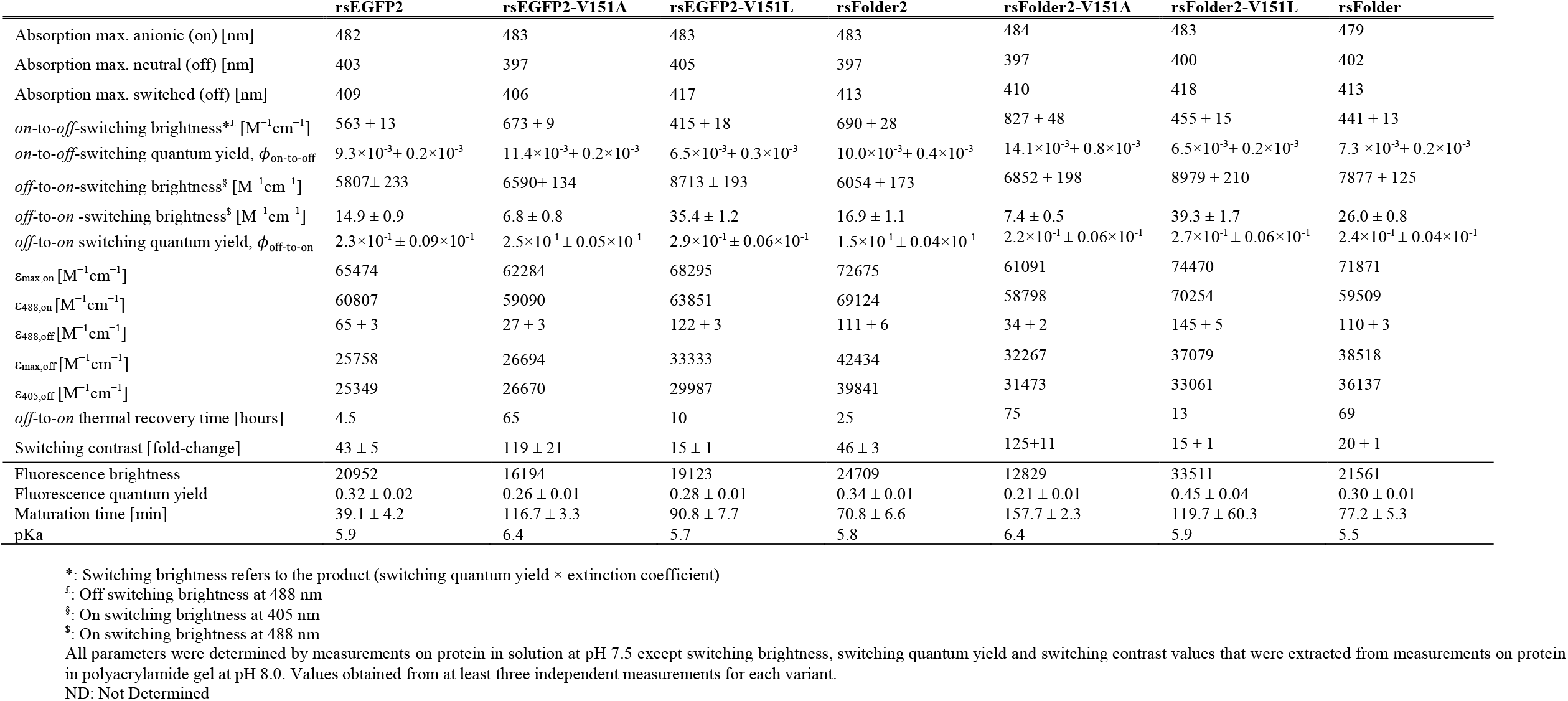
Photophysical parameters obtained *in vitro*

**Figure 3:**
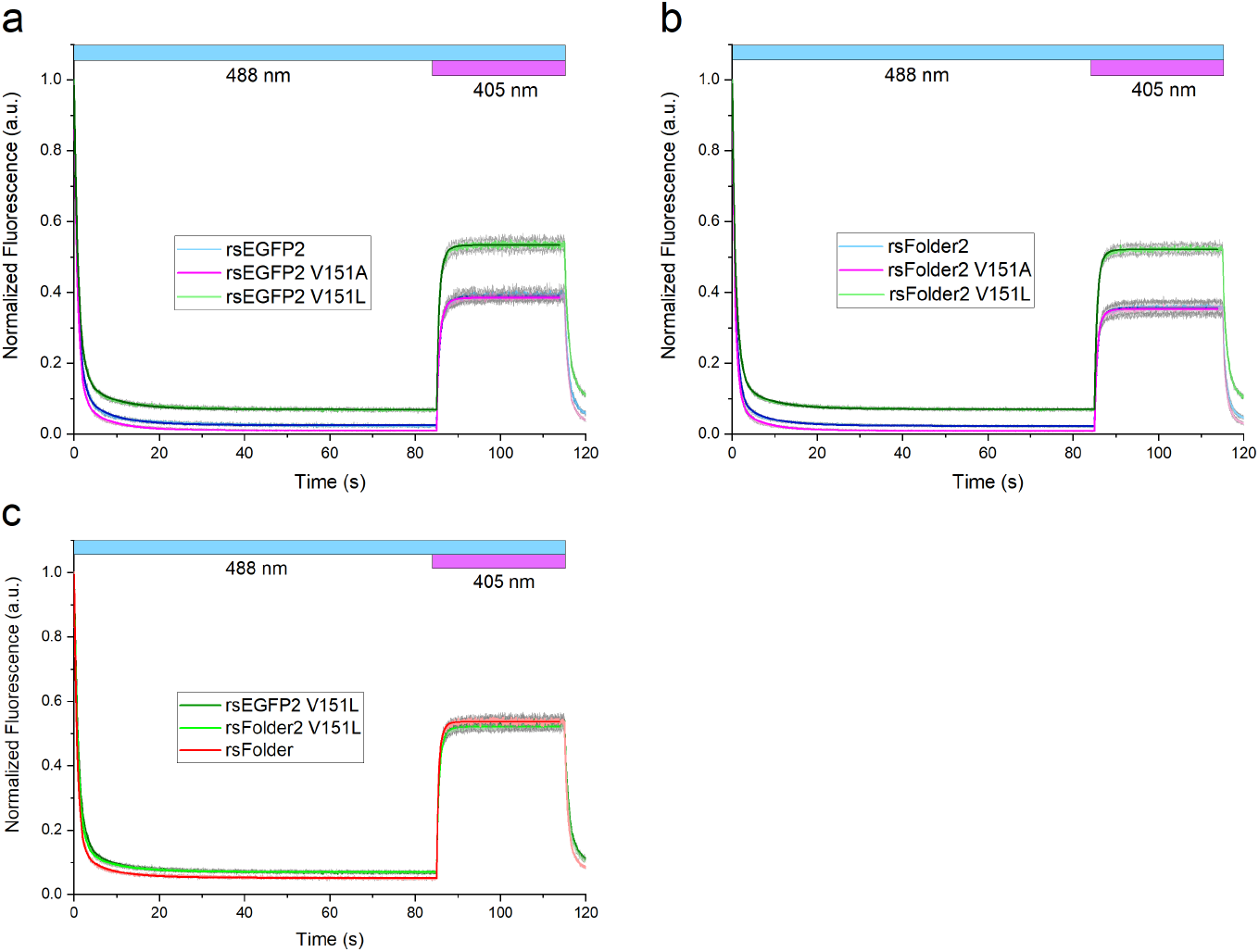
Fluorescence switching curves. (*a, b*) Fluorescence switching curves for rsEGFP2 and rsFolder2 proteins together with their V151A and V151L variants. (*c*) Switching curve for rsFolder compared to V151L variants of rsEGFP2 and rsFolder2. Data obtained from *in-vitro* measurements of purified proteins, embedded in polyacrylamide gel (pH 8.0), on an epifluorescence microscope using 488 nm (0.27 W/cm^2^) illumination throughout data acquisition, and additional 405-nm (0.03 W/cm^2^) during *off*-to-*on* switching. Pale colors stand for the mean values calculated from six measurements with standard deviations shown in grey, whereas dark solid lines represent the fits from the used kinetic model.

To explore the underlying reason for the modified switching contrasts, we examined the photoswitching kinetics of all variants in more detail. Neglecting the slow thermal relaxation in RSFPs (∼hour range) in view of the timescale of our experiments (∼second range), the switching contrast *SC* at wavelength *λ* can be approximated by:

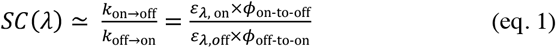

where *k*_on⟶off_ and *k*_off⟶on_ are the *on*-to-*off* and *off*-to-*on* switching rates, *ϕ*_off-to-on_ and *ϕ*_off-to-on_ are the *on*-to-*off* and *off*-to-*on* switching quantum yields, respectively, and ε_λ, on_ and ε_λ, off_ are the extinction coefficients of both the *on* and *off* states at the *off*-switching wavelength, respectively. Changes in the switching contrast can thus be due to changes in switching quantum yields ^[15]^ and/or extinction coefficients of the *on*- and/or *off*-states. For the proteins in the present study, the *off*-switching wavelength is 488 nm. Whereas ε_488,on_ was measured using the Ward method ^[23]^ *ϕ*_on-to-off_, *ϕ*_off-to-on_, ε_488,off_ values were calculated by fitting the experimental fluorescence switching curves (Figure 3) with a kinetic model (see Supplementary methods section and Supplementary text S3). Overall, the dominating effect underlying variations between the observed switching contrasts in the studied variants follows from significant differences in *off*-to-*on* (rather than *on*-to-*off*) switching brightness at 488 nm (Table 1): the V151A variants switch *on* significantly less efficiently at 488 nm than the parent proteins, while the V151L variants switch *on* much more efficiently. The observed differences in the *off*-to-*on* switching brightness are mainly due to differences in ε_488,off_ (reduced ∼2.4 fold in rsEGFP2-V151A, and increased ∼1.9 fold in rsEGFP2-V151L compared to parental rsEGFP2) and only to a minor extend to differences in *off*-to-*on* switching quantum yields (Table 1). Note that absolute values of *off*-to-*on* switching quantum yields are notoriously difficult to determine, as evidenced by the spread in values determined in different laboratories (e.g. for rsEGFP2 in solution, values of 0.12 ^[24]^ and 0.34 ^[21]^ have been published). Yet, values determined under identical conditions in the same laboratory should be comparable. It is thus striking that similar *off-to-on*-switching quantum yields were measured here on rsEGFP2 (0.23) and rsEGFP2-V151A (0.25) embedded in polyacrylamide gels, but a two-fold increase of the quantum yields for the proteins in solution has been reported by us earlier (0.40 and 0.77 for rsEGFP2 and rsEGFP2-V151A, respectively ^[20]^). Unlike data in Table 1, the value for rsEGFP2-V151A in ^[20]^ was determined from a single measurement and we suspect that an unidentified experimental flaw lead to the erroneously high value (0.77) that should thus be discarded. Compared to rsEGFP2 and rsFolder2, the behavior of rsFolder is similar to that of the V151L variants. We note that the V151A variants are characterized by a lower fluorescence brightness than their parents (Table 1).

Overall, our results suggest that the higher and lower switching contrasts of the V151A and V151L variants relative to the parent proteins, respectively, are mainly due to lower absorptions of 488 nm light (ε_488,off_) by the *trans1* chromophore in V151A and to higher absorption of the *trans2* chromophore in V151L.

### UV-visible absorption spectroscopy on rsEGFP2, rsFolder2 and their variants in solution

The differences in ε_488,off_ determined from fitting the experimental fluorescence switching curves (Table 1) can be rationalized by comparing UV-visible absorption spectra of rsEGFP2, rsFolder2 and their V151A and V151L variants in their *on*- and *off*-states, respectively (Figure 4). Indeed, we consistently observed that the maximum of the *off*-state spectra of the V151A variants are blue shifted, and the spectra of the V151L variants red shifted relative to those of the parent proteins (Table 1). In the V151A variants, the blue shifted *off*-state absorbance band leads to a lower residual extinction coefficient at 488 nm (ε_488,off_), resulting in less *on*-state contamination after *off*-switching (Figure 4) and thus to an increased switching contrast. In contrast, the red shifted *off*-state absorbance band in the V151L variants leads to a higher extinction coefficient at 488 nm, more *on*-state contamination and a lower contrast. Interestingly, a shoulder at 440 nm is visible in the *off*-state absorbance spectra of rsEGFP2-V151L, rsFolder2-V151L and rsFolder (Figure 4) that cannot be attributed to residual absorbance of the *on*-state. This shoulder is also visible in *off*-state absorption spectra of microcrystalline rsEGFP2-V151L (Supplementary figure S4b).

**Figure 4:**
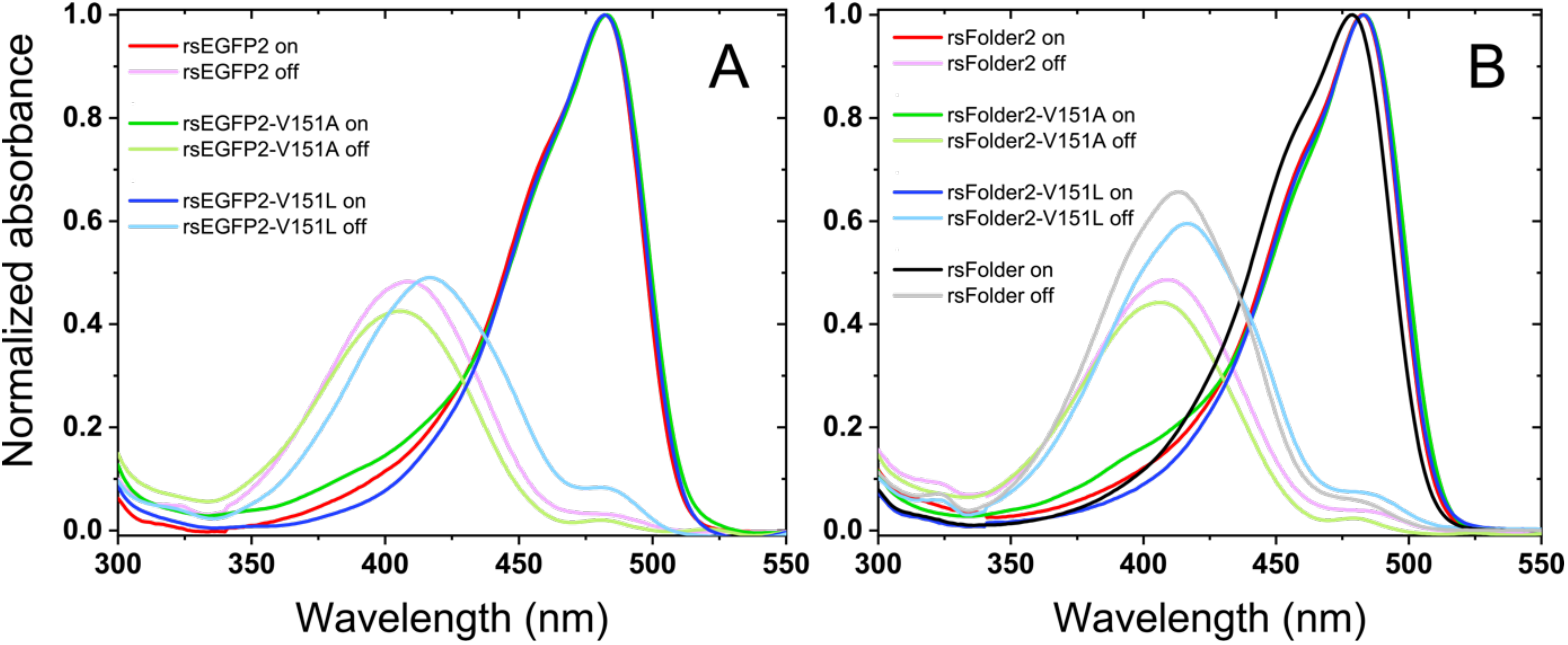
Absorption spectra of fluorescent proteins presented in this study in their fluorescent *on*-state (dark colors) and non-fluorescent *off*-state (dim colors) in solution. (*a*) rsEGFP2 and its variants rsEGFP2-V151A and rsEGFP2-V151L, (*b*) rsFolder2 and its variants rsFolder2-V151A and rsFolder2-V151L and rsFolder. Spectra are normalized relatively to the respective *on* state spectra and are measured in HEPES buffer at pH 7.5.

We also recorded absorption spectra along *on*-to-*off* switching under alternating 488 nm illumination for all variants in the solution state. An isosbestic point (Supplementary figure S7) was always observed, suggesting a homogenous *off*-state, not only for the V151A and V151L variants, but also for the parent proteins. Likewise, all our fluorescent switching curves (Supplementary figure S8) displayed a similar trend, with no sign of more complex kinetics in the parent proteins compared to the V151A and V151L variants.

### Quantum chemical calculation analysis of chromophore conformations in parental rsEGFP2

So far, we implicitly assumed that the *trans1* and *trans2* conformations observed in the crystal structures occur in the proteins in solution, and hence, the blue- and red-shifted absorption could be attributed to *trans1* and *trans2* chromophores, respectively. To test this assumption, the absorption spectra of the *trans1* and *trans2* conformers were characterized by quantum chemistry calculations. Starting from the SFX structure of parental rsEGFP2 (PDB entry 6T39^[18]^), the geometries of models featuring *trans1* and *trans2* chromophore conformations in their respective protein environment (*i*.*e. trans1* being H-bonded to a water molecule and *trans2* to His149; Figure 5a) were optimized. The planarity of the chromophore remained similar to the experimentally derived one (Supplementary table S3) with *trans1* being rather distorted from the planar configuration in contrast to *trans2* (Figure 5a). The two configurations also differ in the length of the phenolic OH bond and bonds of the conjugated system indicating a stronger binding of the phenolic proton concomitant with a reduced π-conjugation in *trans1* as compared to *trans2*. The energy cross sections computed for the phenolic OH bond stretching (Figure 5b) demonstrate a shape typical for the protonated p-hydroxybenzylidene imidazolinone chromophore interacting with a proton acceptor ^[25]^ and confirm a stronger proton binding in *trans1*. Consistent with reduced π-conjugation, the S_0_-S_1_ energy is higher for *trans1* than for *trans2* (Figure 5b, Supplementary table S4), suggesting a blue-shifted absorption maximum for the former. The enhanced proton binging in *trans1* could be linked to a substantial electronic coupling of 15 meV between *trans1* chromophore and electron donor Tyr146. The coupling is facilitated by a H-bond between His149 and Tyr146. The *trans2* chromophore is H-bonded with His149 itself, and its electronic coupling with Tyr146 is reduced to 0.5 meV. The absorption band shapes obtained with a quantum-mechanical model considering the one-dimensional OH-stretching potential (Supplementary figure S9 and Supplementary table S5) also suggest a blue shifted absorption band of *trans1* compared to *trans2*.

**Figure 5:**
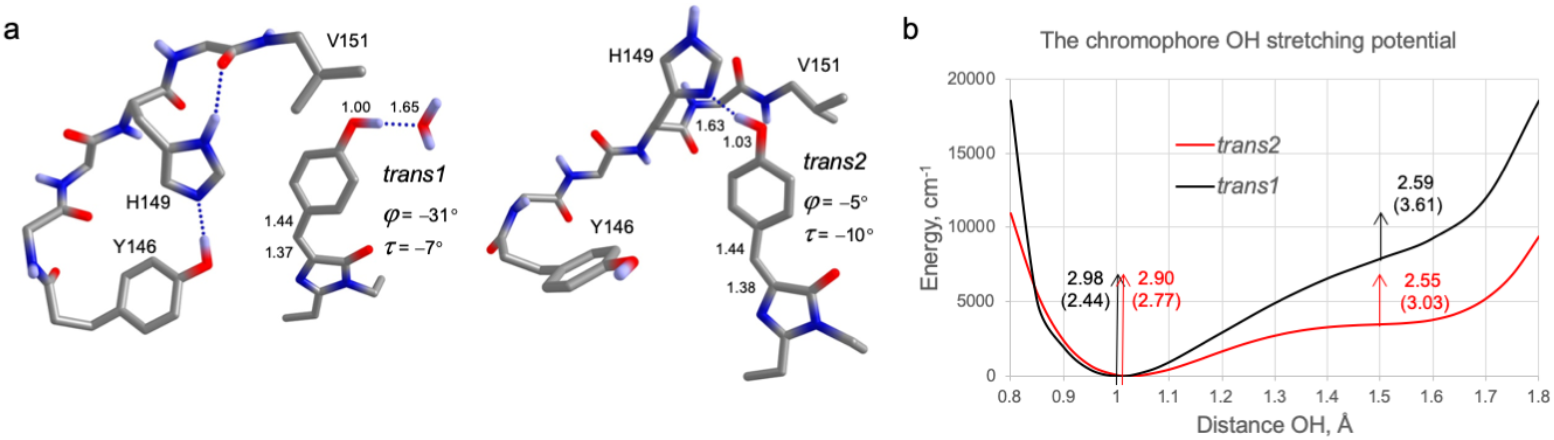
Quantum chemical calculations of *trans1* and *trans2* within parental rsEGFP2. (*a*) Fragments of the chromopore models and selected distances (Å) and angles in the optimized geometries. (*b*) The OH stretching energies. Vertical arrows indicate the S_0_-S_1_ excitation energies (eV) and transition dipole moments (in brackets; Hartree*Bohr). The decrease of the S_0_-S_1_ energy along the OH stretching coordinate is demonstrated. The effect of the OH-stretching potentials on the S_0_-S_1_ vibronic band structure is presented in Supplementary figure S9.

Geometry optimization and excitation energy calculations were performed for the *trans1* and *trans2* models with alanine and leucine, respectively, at position 151. Similar distorted and planar geometries of *trans1* and *trans2*, respectively, were found in these models (Supplementary figure S10 and Supplementary table S3). Further, the calculations confirmed that the S_0_-S_1_ excitation energy is higher for *trans1* compared to *trans2* independent of the residue at position 151 (Supplementary table S4). Hence, our calculations suggest an increase of the excitation energy for *trans1* in comparison to *trans2*, consistent with the blue-shifted absorption band assigned to *trans1*, which we correlate with the increased out of plane distortion and increased proton binding of *trans1*. The calculations thus corroborate our assumption that the *trans1* and *trans2* conformers are adopted both *in crystallo* and in solution.

### Determination of in vivo switching contrasts of rsEGFP2, rsFolder2 and their variants and RESOLFT experiments on rsFolder2-V151A

To investigate the *in vivo* switching properties of *rsEGFP2, rsFolder2 and their variants* at light intensities similar to those typically utilized in RESOLFT nanoscopy, we recorded switching curves on *E. coli* colonies expressing the respective proteins using high light intensities (Figure 6a, b). The determined contrasts of rsEGFP2-V151A (109) and rsFolder2-V151A (119) increased and that of rsEGFP2-V151L (6) decreased with respect to parental rsEGFP2 (28) and rsFolder2 (22), in line with the results obtained with low light intensities *in vitro* (Table 1). We also compared the switching fatigue of parental rsEGFP2 and rsFolder2 and their variants. To this end, the fluorescence of *E. coli* colonies was switched *on* and *off* 4000 times and the maximal fluorescence of the *on*-state was recorded for every switching cycle (Figure 6c). While rsEGFP2 and rsFolder2 could be switched more than 2000 times before their fluorescence was reduced to 50% of the initial brightness, the V151A variants could be switched ∼1500 times. Based on *in vivo* switching properties, both V151A variants appeared to be suitable for RESOLFT nanoscopy.

**Figure 6:**
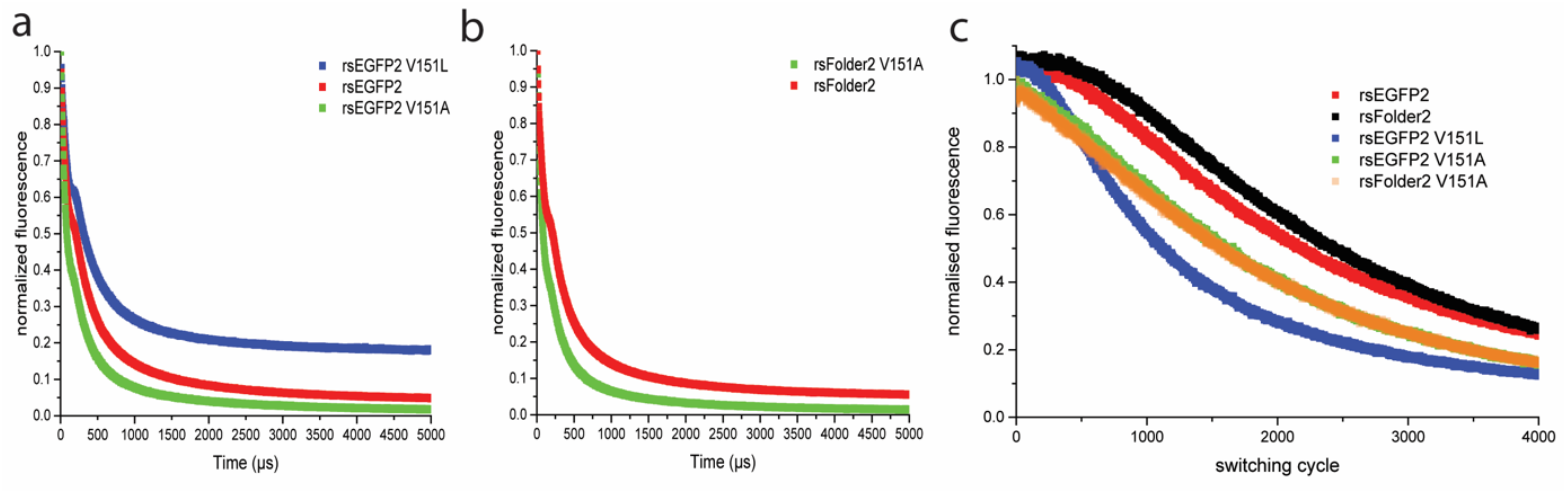
Switching kinetics and switching fatigue of rsEGFP2, rsEGFP2 V151A, rsEGFP2 V151L, rsFolder2 and rsFolder2 V151A. Comparison of the *off* switching curve of rsEGFP2, rsEGFP2-V151A and rsEGFP2-V151L (*a*), as well as rsFolder2 and rsFolder2-V151A (*b*). Switching fatigue measurements of the five proteins (c). All graphs were recorded on living *E. coli* colonies using high light intensities.

For investigating the usefulness for RESOLFT imaging, we decided to concentrate on rsFolder2-V151A, as this variant showed the highest photoswitching contrast *in vitro* (Table 1) and *in vivo*, and could be particularly useful because of its superfolding properties. A fusion protein of the cytoskeletal protein Keratin with rsFolder2-V151A was expressed in cultured human HeLa cells and imaged on the RESOLFT microscope. While the resulting image shows the expected clear improvement in resolution compared to the confocal counterpart (Figure 7), its resolution of ∼60 nm is comparable to the one of previous RESOLFT recordings on the same microscope using parental rsEGFP2 ^[26]^, as judged by the full width at half maximum (FWHM) of line profiles across small Keratin-rsFolder2-V151A filaments. Hence, despite the increased switching contrast of the V151A variant, evidenced *in vitro* (Table 1) and *in vivo* (Figure 6), the recorded images do not show a significant increase in resolution compared to the parent protein. We mainly attribute this result to the lower molecular and cellular brightnesses of these variants compared to the parents (Table 1).

**Figure 7:**
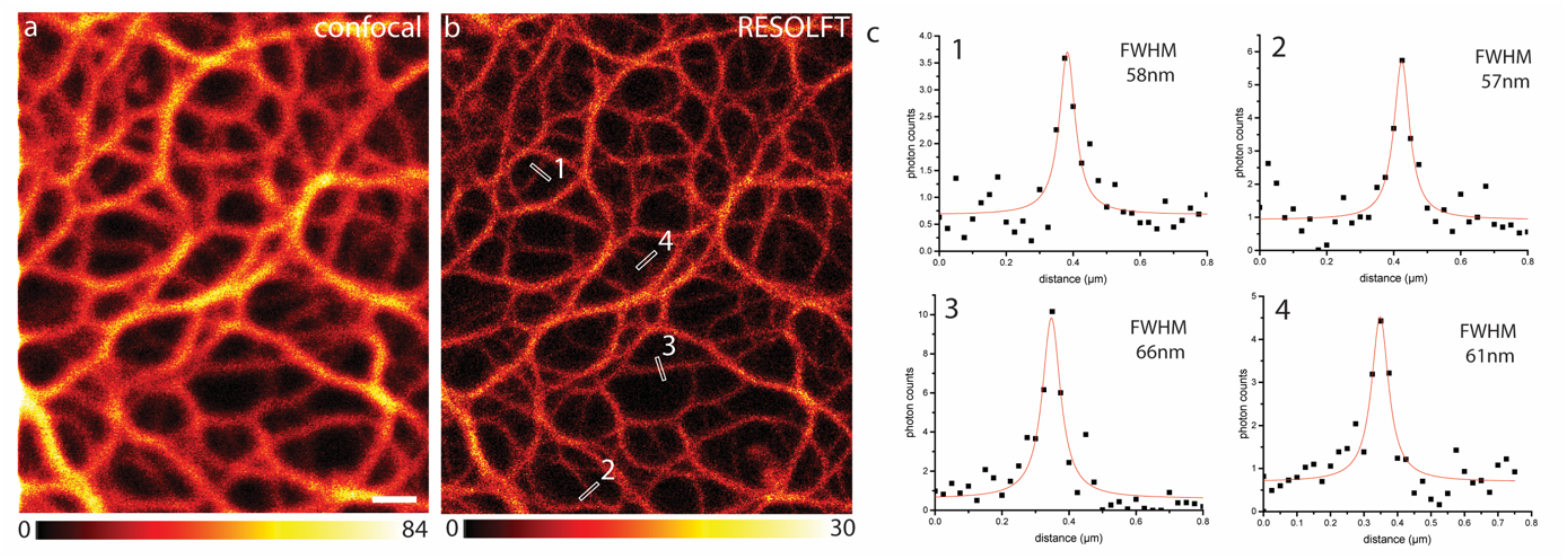
RESOLFT imaging using rsFolder2-V151A. Comparison of a confocal (*a*) and RESOLFT (*b*) recording of a HeLa cell expressing Keratin-rsFolder2-V151A. Line profiles were taken at the indicated positions and the FWHM was determined on the fitted functions (*c*). Scale bar: 1*µ*m. The color maps indicate the actual photon counts.

## Discussion

### Correlation between off-state conformations, switching contrast and absorbance spectra

We establish a link between the switching contrast and *off*-state occupancy of the *trans1* or *trans2* chromophore conformations in rsEGFP2 and rsFolder2 variants by combining our structural and photophysical results. For the *trans1* conformer (V151A variants), the switching contrast is high, whereas the *trans2* conformer (V151L variants) leads to a lower contrast. Parental rsEGFP2 and rsFolder2, exhibiting *trans1/trans2 off*-state heterogeneity (see Supplementary text S4), display an intermediate contrast (Table 1). The quantitative evaluation of fluorescence switching kinetics and the *off-*state absorbance spectra of the investigated variants strongly suggest that the main contribution to modulation of the switching contrast arises from red- and blue-shifted absorption maxima of the *off*-state in the V151L and V151A variants, respectively. This observation correlates with the differences in twist and tilt dihedral angles of the chromophore in *trans1* and *trans2*, as well as in its local environment, that tune the energy gap between the S_0_ and S_1_ electronic states as shown by our high-level quantum chemistry calculations. The red-shift of the experimental absorption maximum of the rsEGFP2-V151L *off*-state was reproduced computationally (Figure 5b) and is consistent with the more extended electron delocalization in the near-planar chromophore of its *trans2* conformation compared to *trans1* (Figure 5a). The less planar *trans1* chromophore conformation might result from stabilization of the protonated form by charge transfer (CT) from Tyr146, activated by H-bonding to His149, that is absent for *trans2*.

When linking structural and spectroscopic results, we rely on the assumption that the *off*-state heterogeneity observed in crystals of parental rsEGFP2 and rsFolder2 is maintained in solution. Our high-level quantum chemistry calculations justify this assumption. Yet, the presence of isosbestic points in the absorbance spectra along *off*-switching (Supplementary figure S7) and detection of a single long-lived *off*-state in our fluorescence-based switching curves (Supplementary figure S11) indicate a homogeneous *off*-state in solution. These findings are consistent if we postulate a fast exchange between *trans1* and *trans2* in parental rsEGFP2 and rsFolder2 in solution, occurring on timescales faster than detectable (∼0.1 s) in our absorbance and switching kinetics measurements. Our measurements in solution thus capture the average photophysical behavior between those of *trans1* and *trans2*, giving rise to an apparently homogeneous *off*-state. The postulated fast exchange contrasts the assumption by Chang et al that *trans* conformations are locked on the second time scale (see Supplementary information S3 in ^[19]^).

### Photoswitching fragility in parental rsEGFP2 and rsFolder2

The differential *trans2* occupancies found in *off*-state structures of rsEGFP2 determined by RT SFX under identical buffer conditions on microcrystals of the same batch but of different age and with varying illumination conditions (Supplementary text S1), as well as the absence of *trans2* in flash-cooled parental rsEGFP2 macrocrystals (Supplementary figure S12) and its presence in rsFolder2 (Supplementary text S2, Supplementary figure S6), suggest that experimental and environmental parameters determine to which extent *trans2* gets populated in parental rsEGFP2 and rsFolder2. We suggest that this is the result of a low, environmentally dependent, barrier in the protein conformational energy landscape ^[27]^ separating access to, or exchange between, *trans1* and *trans2*, a notion that we refer to as “*switching fragility*”. Yet, the experimental conditions that would reproducibly control heterogeneity in the *off*-states of rsEGFP2 and rsFolder2 have not been identified.

Photoswitching fragility in parental rsEGFP2 or rsFolder2 may have consequences for imaging applications, when labels are addressed to various cellular locations with potentially different physicochemical environments. These different environments (e.g. viscosity, ionic strength, or nature of a fusion protein) could lead to different levels of heterogeneity and variability in switching contrast (beyond that expected from pH induced effects ^[11]^). Such variability is expected to be alleviated in rsFolder, the V151L variants, and the high switching contrast V151A variants so that they may be considered as more “robust” than their parents.

In conclusion, this work establishes a causal relationship between the occupancy of two *off* conformations in rsEGFP2 and rsFolder2 and the achievable switching contrast, essentially through absorbance shifts of the *off-*switched chromophore. A point mutation is sufficient to enforce single *trans* conformations in the *off*-state. The rsEGFP2- and rsFolder2-V151A variants, containing only the *trans1* conformer, exhibit greatly enhanced switching contrasts as compared to their parents, both *in vitro* and *in vivo*. Due to a loss in fluorescence brightness of these variants, however, the optical resolution obtained by RESOLFT nanoscopy on Keratin-rsFolder2-V151A filaments did not significantly increase compared to previous studies using either parental rsFolder2 or rsEGFP2. Our rsEGPF2-V151A and rsFolder2-V151A variants constitute promising leads for the next-generation RSFPs, for which the fluorescence brightness has to be increased while maintaining the enhanced switching contrast described here.

## Acknowledgment

We thank Elke De Zitter for critically commenting on the manuscript. The XFEL experiments were carried out at BL2-EH3 of SACLA with the approval of the Japan Synchrotron Radiation Research Institute (JASRI; Proposal No. 2018A8026; 27 – 29 July 2018) and at the CXI beamline at the LCLS (Proposal No. LM47 (23 – 27 June 2016) and LR38 (22 – 26 February 2018). We warmly thank the SACLA and LCLS staff for assistance. Use of the LCLS, SLAC National Accelerator Laboratory, is supported by the U.S. Department of Energy, Office of Science, Office of Basic Energy Sciences under Contract no. DE-AC02-76SF00515. Part of the sample injector used at LCLS for this research was funded by the National Institutes of Health, P41GM103393, formerly P41RR001209. We acknowledge support from the Max Planck Society. The study was supported by travel grants from the CNRS (GoToXFEL) to MW, an ANR grant to MW, MC, MSl (BioXFEL), an ANR grant to DB (grant no. ANR-17-CE11-0047-01), a PhD fellowship from Lille University to LMU and an MENESR – Univ. Grenoble Alpes fellowship to KH, a Russian Science Foundation grant (project #17-13-01051) to TD. This work was partially carried out at the platforms of the Grenoble Instruct-ERIC center (IBS and ISBG; UMS 3518 CNRS-CEA-UGA-EMBL) within the Grenoble Partnership for Structural Biology (PSB). Platform access was supported by FRISBI (ANR-10-INBS-05-02) and GRAL, a project of the University Grenoble Alpes graduate school (Ecoles Universitaires de Recherche) CBH-EUR-GS (ANR-17-EURE-0003). The IBS acknowledges integration into the Interdisciplinary Research Institute of Grenoble (IRIG, CEA).

## Notes

### Competing Interest Statement

The authors have declared no competing interest.

