## Supplementary Information for "Rational control of structural off-state heterogeneity in a photoswitchable fluorescent protein provides switching contrast enhancement"

### Supplementary material

### *Supplementary Methods*

*Site-directed mutagenesis, purification and spectroscopic characterisation of parent proteins (rsEGFP2 and rsFolder2) and their V151A and V151L variants*

The primers for V151A and V151L substitutions were 5′-CAACAGCCACAACGCCTATATCATGGCC-3′ and 5′-CTACAACAGCCACAACCTCTATATCATGGCCG-3′, respectively, for rsEGFP2 and 5′-CAGCCACAACGCGTACATCACC-3′ and 5′-CAGCCACAACCTGTACATCACCG-3′, respectively, for rsFolder2. Mutagenesis, transformation, expression and purification were performed as described in [1]. Absorption spectra were recorded using a Jasco V-630 UV/VIS spectrophotometer (Easton, USA). Emission spectra were measured either with a Jasco FP-8500spectrofluorometer (Easton, USA) or with a CCD-based microspectrophotometer (AvaSpec-ULS2048, Avantes, Eerbeek, The Netherlands) coupled with optic fibres to a cuvette holder and photoinduced fluorescence switching cycles were measured as described earlier [2].

*Microcrystal growth of rsEGFP2 and its V151A and V151L variants, injection for SFX data collection and pre-illumination to induce on-to-off switching*

rsEGFP2, rsEGFP2-V151A and rsEGFP2-V151L microcrystals (3 × 3 × 3 μm$^3$) were generated by the batch method and employing microseeding as described earlier [1,3]. The final protein, precipitant, salt and buffer concentrations were 20 mg/ml, 2 M ammonium sulfate, 20 mM NaCl, 120 mM HEPES pH 8, respectively. Purification and microcrystallization of rsEGFP2, rsEGFP2-V151A and rsEGFP2-V151L took place in April 2015, October 2017 and February 2018, respectively. The rsEGFP2 microcrystal batch was the same as the one used for earlier SFX experiments [1,3]. Prior to injection, sedimented rsEGFP2 microcrystals (but not those of the V151A and V151L variants) were resuspended in 100 mM HEPES pH 8, 2.5 M ammonium sulphate and 5 – 10 % (v/v) of finely crushed rsEGFP2 microcrystals were added to reduce aggregation of microcrystals and the resulting clogging of the injector tubing. All microcrystal suspensions were filtered through a 20-$\mu$m stainless steel filter using a sample loop and a manually-driven syringe. They were loaded into a stainless-steel sample syringe, which was then installed on an anti-settling device [4] onto a Peltier element-cooled (20°C) syringe holder. Crystals were injected at flow rates between 30 – 40 µL min$^{-1}$ with a gas dynamic virtual nozzle (GDVN [5]), using sample capillaries with an inner diameter of 75 µm. For *off*-state data collections, rsEGFP2 microcrystals were injected at a concentration of 20-30% (v/v) into the microfocus vacuum chamber of the CXI instrument [6] at the LCLS (proposal LM47, 23 – 27 June 2016) and those of the V151A and V151L variants at 6.6 % (v/v) into the helium-filled *Diverse Application Platform for hard X-ray Diffraction in SACLA* (DAPHNIS [7]) chamber on the BL2 – EH3 experimental station of SACLA [8](SACLA 2018A8026, 27 – 29 July 2018). For *on*-state data collection of the rsEGFP2-V151A variant, the sample was injected at 5 % (v/v) into the microfocus vacuum back-chamber of the CXI instrument at the LCLS (proposal LR38, 22 – 26 February 2018).

For *off*-state data collections, microcrystals were pre-illuminated during transit from the sample syringe to the injector with 488 nm laser light (200 mW nominal power, 25 ms transit time, 300 W / cm$^2$) within a custom made device [9] to switch them from the resting *on*-state to the *off*-state (Figure 1a). The pre-illumination efficiency, *i.e.* the residual amount of remaining *on*-state was checked by absorption spectroscopy using a Nanodrop 2000c spectrometer. The spectra after 488 nm illumination contain contributions from *off*- and *on*-state chromophores and are distorted by scattering effects. They were modeled by a sum of two Gaussians (Gaussian 1 at 400 nm and Gaussian 2 at 482 nm) and an exponential function, respectively. Baseline distortions due to scattering were subtracted and the spectrum smoothened with a Savitzky-Golay filter and normalized at 280 nm (Supplementary figures S4 and S13). The absorbance of Gaussian 2 at 482 nm relative to the absorbance at 482 nm of the spectrum before 488 nm illumination (assumed to correspond to 100% *on*-state) represents the pre-illumination efficiency. In the case of rsEGFP2-V151A (Supplementary figure S4a) and V151L

(Supplementary Figure S4b), 85% and 77% (accuracy is estimated to be around 10%) were switched to the *off*-state, respectively, and 15% and 23% remained in the *on*-state, respectively. For parental rsEGFP2 (LCLS experiment LM47), only 66% were switched to the *off*-state and 34% remained in the *on*-state (Supplementary figure S13a), *i.e.* less than during our earlier SFX experiments [1, 3], where 90% [1] and 84% (SACLA experiment in 2015 (2015A8031[3]) (Supplementary figure S13b) of parental rsEGFP2 was switched to the *off*-state. We noticed after the LM47 experiment on rsEGFP2 that the SMA fiber within the high-performance liquid chromatography (HPLC) tee union of the pre-illumination device (see Figure 2 in [9]) was dirty or damaged, leading to a 50% drop in laser power at the sample position. For the SACLA 2018A8026 experiment on the V151A and V151L variants, a new version of the pre-illumination device was engineered (Shoeman *et al.*, unpublished).

*RT SFX data collection and online monitoring*

SFX data on parental rsEGFP2 in the *off*-state were acquired with the CSPAD detector [10] operating in a dual-gain mode in the microfocus vacuum chamber of the CXI instrument at the LCLS operating at 120 Hz, with X-ray pulses of 35 fs in length and a photon energy of nominally 9.1 keV (LM47). The X-ray beam was focused to 1 – 2 µm (FWHM). rsEGFP2-V151A *on*-state data (LR38) were collected in the vacuum backchamber of CXI (focus 5 – 10 µm (FWHM); photon energy 9.8 keV). SFX data on rsEGFP2 V151A and V151L variants in their *off*-states were collected on the BL2 – EH3 experimental station (hutch temperature ca 27 - 29°C, but sample syringe on cooled (20°C) Peltier element) using X-ray pulses (nominal photon energy 7.6 keV, pulse length ≤ 10 fs) at a repetition rate of 30 Hz. The X-ray beam was focused to 1.3 µm (H) × 1.4 µm (V) (FWHM), the pulse energy was ~400 µJ. Data were acquired with the octal-MPCCD detector with eight sensor modules positioned 50 mm away from the sample. The CFEL–ASG Software Suite (CASS [11]) was used for online monitoring of diffraction data, such as diffraction resolution, hit rate, fraction of multiple hits and pixel saturation.

*SFX data processing*

NanoPeakCell [12] was used for the on- and off-line hit finding. Hit-finding parameters were adjusted after visual inspection of the first diffraction patterns using the NanoPeakCell graphical interface.

For rsEGFP2, rsEGFP2-V151A in the *on-* and the *off*-state, and rsEGFP2-V151L in the *off*-state, 80 605, 17 538, 10 794 and 4930 images were indexed, respectively (Table S2). The CrystFEL suite [13] was used for further data processing. The data were indexed with *Mosflm* and integrated using v.0.6.2 for parent rsEGFP2, v.0.6.3 for rsEGFP2-V151A in the *on*-state and v.0.7.0 for rsEGFP-V151A and rsEGFP2-V151L in the *off*-state. Merging was performed using CrystFEL v.0.8.0 with the Monte Carlo algorithm, *process_hkl*, for the parental rsEGFP2 *off*-state dataset and with *partialator* using the xsphere partiality model with one cycle of scaling and post-refinement for all rsEGFP2-variant datasets. A resolution cut-off of 1.7, 1.9, 1.95, 2.1 Å was chosen based on SNR, Rsplit and CC* statistics (Table S2) for parent rsEGFP2, rsEGFP2-V151A in the *on*-state, and rsEGFP2-V151A and rsEGFP2-V151L in the *off*-state, respectively.

*SFX structure solution and refinement*

The SFX structures were phased by molecular replacement with *Phaser* [14]. As a search model was used the SFX *off*-state structure of the parental rsEGFP2 (PDB entry 6T39 [3]) for structures of rsEGFP2 in the *off*-state (PDB entry 7O7U), and of rsEGFP2-V151A (PDB entry 7O7X) and -V151L (PDB entry 7O7W) in the *off*-state. The SFX parental rsEGFP2 *on*-state structure (PDB entry 5O89[11]) was used as a search model for the structure of rsEGFP2-V151A in the *on*-state (PDB entry 7O7V).

Refinement included positional and isotropic individual B-factor refinement in reciprocal space in Refmac5 [15] for all structures. Model building and real space refinement were performed in *Coot* [16]. Initially, the chromophore had been omitted from the model and subsequently included and modelled as described below. Once the chromophore had been included, its position and isotropic individual B-factors were refined in reciprocal space, while the protein moiety remained fixed. All the structures were superposed to the SFX parental rsEGFP2 *off*-state structure (PDB entry 6T39 [3]) using the CCP4 program *superpose* which is a structural alignment tool based on secondary structure matching.

For rsEGFP2 in the *off*-state, the chromophores were modelled in a *trans1* conformation at 70% occupancy and the *on*-state *cis* conformation at 30% according to spectroscopy (Supplementary figure S13a). The resulting $F_{obs} - F_{calc}$ map showed a positive peak indicating the presence of *trans2* (Supplementary figure S2a). When the occupancy of *cis*, *trans2* and *trans1* were set to 30%, 30% and 40% occupancy the resulting $F_{obs} - F_{calc}$ map was featureless in the chromophore region (Supplementary figure S2b). That a considerable fraction (30%) of the chromophore was in the *trans2* conformation was also confirmed by a $F_{obs} - F_{calc}$ map in which the hydroxybenzylidene moiety was omitted (Supplementary figure S2c). We note that the chromophore occupancies were different from the ones in parental rsEGFP2 in the *off*-state reported earlier (6T39 [3], 10% *cis*, 25% *trans2* and 65% *trans1*), in agreement with spectroscopy (Supplementary figure S13).

For rsEGFP2-V151A in the *on*-state, the *cis* chromophore of parent rsEGFP2 was included at 100% occupancy.

For rsEGFP2-V151A in the *off*-state the chromophore was first modelled at 100% occupancy in the *trans1* conformation of the parent rsEGFP2 (PDB entry 7O7U) resulting in negative $F_{obs} - F_{calc}$ peaks on the chromophore and the side chains of Tyr146 and H149 (not shown). Then, the chromophore (*cis*) and Tyr146 and H149 conformations of the rsEGFP2-V151A SFX *off*-state model (PDB entry 7O7X) were added and their occupancies set manually. Several cycles of positional and isotropic individual B-factor refinements in reciprocal space were carried out. No major features remained in the resulting $F_{obs} - F_{calc}$ map when the *cis* and *trans1* chromophores were occupied at 20 and 80%, respectively (Supplementary figure S3b), in accordance we spectroscopy (Supplementary figure S4a) that indicated 85% of the molecules were switched to the *off*-state.

For rsEGFP2-V151L in the *off*-state the *trans2* chromophore of parental rsEGFP2 (PDB entry 7O7U) and the *cis* chromophore of the rsEGFP2-V151L cryo-MX *on*-state model (PDB entry 7O7E) were inserted and their respective occupancies varied manually. The resulting $F_{obs} - F_{calc}$ map was featureless when the *cis* and *trans2* chromophores were occupied at 25 and 75%, respectably (Supplementary figure S3c), in accordance we spectroscopy (Supplementary figure S4b) that indicated 77% of the molecules were switched to the *off*-state.

*Crystallization of rsEGFP2-V151A and -V151L for synchrotron cryo-crystallography*

For V151L, crystallization was carried out by vapor diffusion with hanging drops in 24-well plates. The well solution was composed of 100 mM HEPES pH 7.8 - 8.4 and 1.7 - 2.4 M ammonium sulfate for a total volume of 500 µl. Drops were prepared at 20 °C by mixing 2 µl from the well with 2 µl of protein solution at 10 mg/ml in 50 mM HEPES pH 7.5, 50 mM NaCl. Crystals appeared within a few days and continued growing during a few weeks to reach their final size.

For V151A, crystallization trials were first assessed following the same procedure using the same conditions than for V151L but without success (drops remained clear even after several days). A cross-seeding approach was then adopted. First, V151A crystals were generated by vapor diffusion in sitting drops on 24-well plates using seeds of parental rsEGFP2. The well solution was composed of 100 mM HEPES pH 8, 2 M ammonium sulfate. Drops were prepared at 20 °C by mixing 10 $\mu$l of protein solution at 10 mg/ml, 50 mM HEPES pH 7.5, 50 mM NaCl with 10 $\mu$l of solution at 2 M ammonium sulfate, 100 mM HEPES pH 8 containing parental rsEGFP2 seeds diluted 1/1000 (v/v). Crystals appeared after a few days and were used for generating seeds that were used to crystallize V151A on 24-well plates using the vapor diffusion method in hanging drops. The well solution was composed of 100 mM HEPES pH 8, and 1.8 - 2.4 M ammonium sulfate concentration. Drops were prepared by mixing 2 $\mu$l of protein solution at 10 mg/ml, 50 mM HEPES pH 7.5, 50 mM NaCl with 2 $\mu$l containing V151A seeds in 2 M ammonium sulfate, 100 mM HEPES pH 8. Crystal appeared and reached their final size within a few days. Generation of V151A macrocrystals without seeding was achieved only after V151A structures had already been determined.

*In crystallo photoswitching, synchrotron data collection and processing of rsEGFP2-V151A and -V151L variants*

For structure determination of the rsEGFP2-V151A variant in its *on*-state, a macrocrystal (200 × 20 × 20 $\mu m^3$) grown in 2.2 M ammonium sulfate, 100 mM HEPES pH 8 was harvested with a cryo-loop and soaked for a few seconds in a cryo-solution composed of 2 M ammonium sulfate, 100 mM HEPES pH 8, 20 % glycerol (the same cryo-solution was used for all crystal preparations presented here). Then the crystal was flash-cooled in liquid nitrogen. For structure determination of V151A in the *off*-state a macrocrystal (150 × 10 × 10 $\mu m^3$) grown in the same condition was exposed to 488 nm laser-light, guided through a fiber, for around one minute at room temperature (RT) directly in the drop where it grew until the crystal turned colorless. The crystal was then harvested with a cryo-loop and soaked in the cryo-solution before being flash-cooled. For structure determination of the rsEGFP2 V151L variant in its *on*-state, a macrocrystal (300 × 100 × 100 $\mu m^3$) grown in 1.7 M ammonium sulfate, 100 mM HEPES pH 7.8, 10 mg/ml was harvested with a cryo-loop and soaked in the cryo-solution before being flash-cooled in liquid nitrogen. For structure determination of V151L in the *off*-state, the same procedure than for V151A was applied. A macrocrystal (150 × 50 × 50 $\mu m^3$) grown in 1.8 M ammonium sulfate, 100 mM HEPES pH 8.2, 10 mg/ml protein was exposed to 488 nm laser-light for around one minute at RT as described above and then soaked in the cryo-solution before being flash-cooled.

X-ray crystallographic oscillation data were collected at 100 K on ID29 [17] at the European Synchrotron Radiation Facility (ESRF, Grenoble, France) operating in 7/8 multibunch mode at an X-ray wavelength of 0.976 Å (12.7 keV) and with a beamsize of 30 $\mu$m × 30 $\mu$m. Data were recorded with a Pilatus 6M detector (Dectris). For rsEGFP2-V151A in the *on*-state, 1050 frames were recorded with an oscillation range of 0.10 ° and an exposure time of 0.037 s per frame with a photon flux of 1.6 × $10^{11}$ ph/s at 5% transmission. For rsEGFP2-V151A in the *off*-state, 547 frames were recorded (oscillation range 0.15 °, exposure time 0.039 s per frame, photon flux 2.8 × $10^{11}$ ph/s, 9.94 % transmission). For rsEGFP2 V151L in the *on*-state, 794 frames were recorded (oscillation range 0.15 °, exposure time 0.1 s per frame, photon flux 2.2 × $10^{10}$ ph/s, 1.27 % transmission). For rsEGFP2 V151L in the *off*-state, 950 frames were recorded (oscillation range 0.10 °, exposure time 0.037 s per frame, photon flux 2.8 × $10^{10}$ ph/s, 9% transmission). Images were indexed and intensities were integrated and merged using XDS [18].

*Molecular replacement and structure refinement of rsEGFP2-V151A and -V151L synchrotron cryo-data*

For both states of the two variants, molecular replacement was performed with *Phaser* [14] using the cryo-structure of rsEGFP2 in the fluorescent *on*-state as a starting model (PDB entry: 5DTX [19]). Manual model building was performed in *Coot* [16] and refinement with the *Phenix* suite [20] included positional and isotropic individual B factor refinement. *PyMOL* [21] was used to produce figures. For the structure of the *on*-state of rsEGFP2-V151A (PDB entry 7O7D), the occupancy of the phenol of the chromophore and the imidazole of H149 was fixed to 0.9 before final cycles of refinement were carried out. For the *off*-state structure of rsEGFP2 V151L (PDB entry 7O7H), the chromophore, H149 and L151 were modeled in alternate conformations.

We note that *on*-state crystal structures of the rsEGFP2 V151A variant have been determined by both synchrotron cryo-crystallography (PDB entry 7O7D, Supplementary table S1; Supplementary figure S5a) and RT SFX (PDB entry 7O7V, Supplementary figure S3a) and the one of the V151L variant only by synchrotron cryo-crystallography (PDB entry 7O7E, Supplementary table S1, Supplementary figure S5c).

*Crystallization, synchrotron cryo-crystallographic data collection and structure solution and refinement of rsFolder2 in its on- and off-states* rsFolder2 was crystallized at a concentration of ~12 mg/ml using the vapor diffusion method, against a reservoir consisting in 20% PEG 3350, 100 mM Tris pH 8.5 and 20 mM NaCl. Thick plate-shaped crystals appeared after 3-4 days at 20 ºC. Before harvesting, crystals were cryo-protected using mother liquor solution supplemented with 15% glycerol and then cryo-cooled in liquid nitrogen. To obtain the *off*-state structure, crystals were illuminated for 5 s with an optical fiber connected to a 488 nm laser just before harvesting. X-ray diffraction data collection was performed at beamline ID30a-3 (MASSIF-3) of the ESRF at 100K, using 0.97 Å wavelength X-rays and a Pilatus3 2M detector (Dectris Ltd.). Data processing was carried out with XDS [18], and merged data were phased by molecular replacement with Phaser [14], within the CCP4i2 suite [22], using as search model the rsFolder *on*-state structure (PDB entry 5DTZ[19]). Final models were obtained after successive refinement rounds (maintaining isotropic B-factors) with Refmac5 [15], with manual model building in-between cycles using *Coot* [16]. For the *on*-state structure (PDB entry 7AMB), a fully occupied *cis* chromophore was found in the active site, as expected. In the case of the *off*-state structure (PDB entry 7AMF), initially only the *cis* and *trans1* conformers were modelled. After a round of refinement, there was a faint indication of the presence of the *trans2* conformer as a weak positive peak in the $F_{obs} – F_{calc}$ map. This evidence was clearer when the NCS averaged difference map (four monomers in the asymmetric unit) was calculated with *Coot* (Supplementary figure S6 b). The *trans2* conformation was modelled in each of the monomers and then refined together with the *cis* and *trans1* conformations. Occupancies for the different conformers were set manually according to electron density in the $2F_{obs} – F_{calc}$ and $F_{obs} – F_{calc}$ maps, and then assessed after refinement to make sure there were no further peaks in the $F_{obs} – F_{calc}$ map. Validation of the structures was performed using Molprobity [23], JCSG QC-check server, and the validation tools within *Coot* as well as the PDB validation service.

*Crystallization, in crystallo photoswitching at various illumination intensities, synchrotron data collection and processing of parental rsEGFP2*

Crystals of parental rsEGFP2 were prepared as previously described [19]. Crystals were then individualized within crystallization trays and exposed to controlled power densities of 488 nm laser light (0.1, 0.2, 0.4 and 0.8 W/cm$^2$) during 30 s. Crystals were then immediately cryoprotected in the mother liquor solution supplemented by 15% glycerol, mounted on microloops and flash-cooled in liquid nitrogen. X-ray data collections was performed at the ID29 beamline at the European Synchrotron Radiation Facility (ESRF, Grenoble, France) operating in 7/8 multibunch mode at an X-ray wavelength of 0.976 Å (12.7 keV), with a beamsize of 30 $\mu$m × 30 $\mu$m and recorded with a Pilatus 6M detector (Dectris).

*Fluorescence microscopy*

Ensemble fluorescence microscopy data were acquired on a custom-made setup based on an Olympus IX81 inverted microscope equipped with a x100 1.49 NA oil-immersion apochromatic objective lens (Olympus, Japan). Widefield illumination was achieved by focusing the diode-pumped solid state 405-nm (CrystaLaser, USA) and 488 nm (Spectra Physics, USA) laser beams to the back focal plane of the objective. Intensities of laser illuminations at the sample were tuned by an acousto-optical tunable filter (AOTF, AA Opto Electronic, France). Fluorescence images were acquired with an Evolve 512 back-illuminated EMCCD camera (Photometrics, USA) controlled by the Metamorph software (Molecular Devices, USA).

*Fluorescent-protein (FP) immobilization on a polyacrylamide gel*

5 $\mu$L of purified FP at ~12 mg/ml concentration was mixed with 50 $\mu$L of 30% polyacrylamide (PAA) solution with 43 $\mu$L of Tris at pH 8.0. PAA polymerization was initiated by adding 1 $\mu$L of 10% APS (ammonium persulfate) and 1 $\mu$L TEMED (Tetramethylethylenediamine). Twenty five $\mu$L of the FP-containing PAA was then spread between two glass coverslides separated by a calibrated spacer (Gene Frame 25 $\mu$l (1.0 × 1.0 cm) AB-0576 Thermo Scientific) thoroughly cleaned using a UV ozone cleaning system (HELIOS 500, UVOTECH Systems) to form a uniform 200 $\mu$m thick layer, and let to harden at room temperature for 5 minutes.

*Ensemble-level switching experiments*

Ensemble level switching experiments were based on measuring the evolution of fluorescence emission. The illumination scheme for driving *off*- and *on*-switching in all variants was the following: first, samples were submitted to 0.086 W/cm$^2$ of 405 nm light for 2.5 s, so as to maximize the population of proteins in the *on*-state. *Off*-switching was performed by recording 1700 EMCCD frames of 25 ms exposure at 20 Hz with dark periods of 25 ms in between frames. During frame exposure, actinic light at 488 nm (0.27 W/cm$^2$) also served as excitation light to readout fluorescence. *On*-switching was performed by recording 600 frames using the same scheme, except that the dark periods of 25 ms in between frames were replaced by illumination at 405 nm (0.03 W/cm$^2$). Thus, FPs were first *off*-switched for 85 s under 488 nm light, and subsequently switched-*on* by 405 nm light, in the presence of 488 nm light, for 30 s. The laser power at the sample position was measured with a calibrated power meter (S170C sensor and PM100D console, Thorlabs), and laser power densities were estimated from a Gaussian fit of the laser profiles that were recorded with fluorescent coverslides (92001, Autofluorescent Plastic Slides, Chroma). Each variant was measured three times from at least three independently prepared samples.

*Extraction of switching curves*

Analysis of switching cycles was performed on a small region of 5 × 5 pixels (0.38 $\mu m^2$) located at the region of maximum laser intensity, so as to ensure homogenous illumination. Of note, considering the thickness of the FP samples (200 $\mu$m), which exceeds the depth of field of the employed X100 objective (~500 nm), the signal analyzed in this small region unavoidably contained residual contributions from out-of-focus FPs submitted to lower illumination intensities. This out-of-focus effects were corrected by estimating the contribution of FPs located elsewhere than in the selected region. This contribution was based on estimating the depth-dependent point spread function (PSF) of the microscope objective according to a Gibson & Lanni model using the PSF generator plug-in of Fiji version 1.52p.

*Determination of switching quantum yields and extinction coefficients $\varepsilon_{488,off}$*

*On*-to-*off* and *off*-to-*on* photoswitching parameters were derived from global fitting of both *off*-switching and *on*-switching phases of the switching cycles with a kinetic model that includes, in addition to the *on*- and *off*-states, a short-lived reversible dark states that can be accessed from the *on*-state and that needs to be invoked to account for the strongly biphasic *off*-switching decay curves. Due to the strong photofatigue resistance of all tested variants, photobleaching was assumed to not contribute to the first photoswitching cycle studied here. The fitting model also took into account the effect of switching during the integration time of the EMCCD, which can be non-negligible for the fast switching proteins studied here. Using measured laser power densities and molar extinction coefficients of the *on*-state proteins obtained by the Ward method (see below) as input, the global fitting of both the *on*-to-*off* and the *off*-to-*on* phases of the switching cycles allowed retrieving, in addition to the *on*-to-*off* and *off*-to-*on* switching quantum yields, the extinction coefficient of the *off*-state at 488 nm, which largely dictates the experimentally observed switching contrast. Fitting was realized with a custom-made routine based on the lsqcurvefit method in Matlab.

*Measurement of absorption spectra before and during photoswitching*

Purified proteins of rsEGFP2, rsFolder2 and their related variants V151A and V151L were adjusted in concentration to ~0.5 OD at their maximum anionic peak absorption in a 0.1 M HEPES buffer, pH 7.5. These samples were placed in a quartz cuvette at room temperature in a CCD-based microspectrophotometer [2] equipped with a square diffuser within the excitation path to achieve homogeneous illumination. Before acquiring each spectrum, the sample was pre-exposed to 405 nm laser light for ~5 s (~88 mW/cm$^2$) to maximize the *on*-state. Photoswitching data was obtained by alternating laser illuminations at 488 nm (~25 mW, 730 ms) and synchronized spectrometer acquisition with white lamp probing (70 ms).

*Determination of off-to-on thermal recovery time*

After switching the RSFPs to their off state by constant illumination at 488 nm, their thermal relaxation was probed by monitoring the absorption as a function of time for 1 to 5 days. Diluted proteins (50 µl) were placed at 20°C in a 50-µl cuvette within a CCD-based spectrometer (AvaSpec-ULS2048, Avantes, Eerbeek, The Netherlands) coupled with optic fibres to a cuvette holder. Absorption spectra were measured every 10 minutes (or 20 minutes for slower mutants) thanks to a pulse of white light synchronised to the spectrophotometer acquisition.

*Determination of maturation time*

Fluorescent proteins were overexpressed in *E. coli* BL21(DE3), transformed to express fluorescent proteins and plated. Three colonies for each FP were transferred in individual flasks containing 100 mL LB medium and allowed to grow for 4 hours before IPTG-induction. The agitation (230 rpm, 37°C) was further pursued for 1 hour to allow the overexpression of FPs and to ensure the exponential growth log phase of bacteria. Protein synthesis was then stopped by the addition of chloramphenicol (300 µg/mL) and 100 µL of each culture was deposited within a 96-cell plate. Absorption and emission points were recorded every 5 min using a Biotek Synergy H4 microplate reader (Winooski, VT, USA) under agitation and controlled temperature at 37°C. Absorption at 600 nm was used to normalize the emission values that were obtained using excitation at 485±20 nm and emission and 528 ±20 nm. Maturation time was evaluated by fitting a logarithmic model and determining the half-time of each fluorescence increase.

*Measurement of switching kinetics and switching fatigue in vivo*

The measurements of switching kinetics and switching fatigue under conditions comparable to RESOLFT nanoscopy conditions were performed on *E. coli* colonies using an automated microscope. Fluorescent proteins were expressed in bacteria grown on LB agar plates containing 0.02% L-Arabinose for 20 h at 37°C using the pBAD expression system and *E. coli* Top10 cells. The *E. coli* colonies were irradiated with laser light of 405 nm for 2 ms (to switch RSFPs completely to the *on*-state) and of 488 nm for 5-20 ms (to switch RSFPs completely to the *off*-state). A 20-fold objective (numerical aperture (NA) 0.4) was used to focus light onto the colonies with power of 5 mW (488 nm) and 1.5 mW (405 nm). The fluorescence was detected with a photomultiplier tube. For measurement of the switching kinetics, the proteins were switched *on* and *off* ten times consecutively, the average of the ten *off* switching curves was calculated for every colony and then normalized to the initial colony brightness. The graphs in figure 6 represent the average of 20 colonies, respectively. For quantification of switching fatigue, the colonies were switched *on* and *off* completely for 4000 times consecutively and the fluorescence intensity of the first time point of every *off*-switching step was used for quantification of the colony brightness in the respective switching cycle. The data was normalized to the initial colony brightness. The graphs represent the average of 10 colonies, respectively.

*RESOLFT microscopy*

HeLa cells were transfected with the plasmid pKeratin-rsFolder2-V151A using TurboFect™ according to the manufactures' instructions. Imaging was performed 20 h after transfection in DMEM without phenol red (Thermo Fisher Scientific, Waltham, MA, USA) at room temperature. RESOLFT microscopy was performed using a customized 1C RESOLFT QUAD scanning microscope (Abberior Instruments, Göttingen, Germany) equipped with a 100X oil immersion objective (NA 1.4). The RESOLFT image was recorded by applying the following three illumination steps at each scanning position. First, proteins were transferred to the *on*-state by illuminating with 405-nm light for 15 µs with an power of 2 µW. Second, the proteins in the periphery of the focal spot were transferred to the *off*-state using a doughnut-shaped beam of 488 nm light with an intensity of 15 µW for 520 µs. Finally, residual *on*-state proteins at the centre of the spot were excited for 30 µs with 8 µW 488 nm light. In the last step, fluorescence was detected using a SPCM-AQRH-13 photon counting module (Excelitas

Technologies, Waltham, MA, USA) with a HC 550/88 detection filter. Power values were measured in the back focal aperture of the objective lens. The scanning step size was set to 25 nm. A line accumulation of 2 was used. The corresponding confocal image was taken accordingly without the *off*-switching step. The full width half maximum (FWHM) of keratin filaments (Figure 7 was determined by averaging three adjacent line profiles at the indicated positions in the raw data image and fitting with a Lorentz curve using Origin 9.1.

*Quantum chemical calculations*

The Firefly program version 8.0 (http://classic.chem.msu.su/gran/firefly/index.html), which is partially based on the US GAMESS code[24], was used to perform quantum-chemistry calculations. Geometry optimization and OH-stretching energy cross sections were computed with the B3LYP-D3 method[25]. The excitation spectrum and electronic couplings were computed with the XMCQDPT2 method [26]. To estimate the $S_0$-$S_1$ energy and transition moments, the SA3-CASSCF(2e,2MO) zero-order wave function was used (two electrons in two molecular orbitals of the chromophore were correlated and state averaging was performed for three states), whereas to compute electronic coupling using the generalized Mulliken-Hush method [27] the SA4-CASSCF(4e,3MO) zero-order wave function (the active space was extended to four electrons in three molecular orbitals by additionally including the occupied orbital of Tyr146; four states were included in state averaging) was used in the XMCQDPT2 calculations. The cc-pvdz basis set [28] was used throughout. The geometries of the cluster models were optimized starting from the coordinates PBD 6T39[3]. To model the active sites containing *trans1* and *trans2* chromophore geometries the subsets of coordinates labelled as A and C, respectively, were selected from the PBD file. Fig. 5 presents the results of the rsEGFP2 active-site models comprised of 283 atoms. The cluster models included the following fragments of the chromophore-containing site: Thr63-Leu70, Gln95, Arg97, Tyr146-Tyr152, Ser164, Gln184, Asn186, Thr204-Ser206, Glu223 and water molecules. The ionisable groups were set as follows: Arg97 protonated, chromophore neutral, Glu223 neutral, His149 neutral (δN-H for *trans1* and εN-H for *trans2*), hence, the cluster model has a net charge +1. During geometry optimization, the coordinates of selected atoms from the cluster boundary were kept constant. The OH stretching potentials of the chromophore phenolic group were computed starting from the optimized geometries by changing the coordinates of the H atom, whilst all other coordinates were kept as in the optimized structure. At the optimized geometries and geometries with the phenolic OH distance 1.5 Å, excitation energies were computed. To model the band shape of the $S_0$-$S_1$ transition, active-site models of a reduced size were employed (for details see caption of Supplementary figure S9). To obtain the A151 and L151 models, the V151 side chain was modified accordingly, and similar computations as for the rsEGFP2 cluster models were performed.

### *Supplementary Text*

*Supplementary text S1: Occupancies of trans1 and trans2 off-state chromophore conformations in parental rsEGFP2 determined from data collected during three different SFX experiments*

Occupancies of chromophore conformations differ in *off*-state structures determined from data collected during three different SFX experiments, *i.e.* 10%, 90% and 0% [1], 10%, 65% and 25% [3], 20%, 40% and 40% (present work) for *cis*, *trans1* and *trans2*, respectively (Supplementary figure S2). The conversion efficiency differed in the latter two SFX experiments as assessed by absorption spectroscopy (Supplementary figure S13), with 10% [3] and 20% (present work) of the molecules remaining in the *on*-state, suggesting that illumination conditions may modulate the *trans1*/*trans2* ratio. Alternatively, aging of the crystalline proteins might be at the origin of *trans2* being increasingly populated in *off*-state structures of rsEGFP2 determined by SFX under identical buffer conditions on microcrystals of the same batch (0% in May 2015 [1], 25% in July 2015 [3] and 40% in June 2016 (present work)).

*Supplementary text S2: Synchrotron cryo-crystallography of parental rsEGFP2 crystals following RT illumination*

To study conformational *off*-state heterogeneity by synchrotron cryo-crystallography, macrocrystals of parental rsEGFP2 in their *on*-state were illuminated with 488 nm light at RT at various intensities for 30 s, flash-cooled and the resulting structures determined (Supplementary figure S12). With increasing illumination intensity, the occupancy of the *trans1* chromophore conformation increased at the expense of the *cis* conformation, yet the *trans2* conformation was never observed. This suggests that *trans2* does not build up to the same extent in rsEGFP2 macrocrystals, or is eliminated upon flash cooling.

*Supplementary text S3: Analysis of photoswitching curves*

We first observed that *on*-to-*off* switching of all investigated variants exhibits a biphasic behavior (Supplementary figure S8), suggesting the transient build-up of a short-lived dark state of potentially different spectroscopic and/or structural nature as compared to the *off*-states associated with *trans1* and *trans2* investigated in the present study. Such additional dark states were previously observed in other RSFPs and in photoconvertible FPs [29], and may involve a twisted *cis* configuration of the chromophore with rapid thermally induced relaxation, but their exact nature will need to be investigated in future studies.

The V151A variants switch *off* slightly more efficiently upon illumination at 488 nm than the parent proteins (*off*-switching rate increased by +20%), and the V151L variants less efficiently (-30%). This appears to be mostly an effect of moderate differences in *off*-switching quantum yields. Upon illumination at 405 nm, the V151A variants switch *on* about as efficiently as the parent proteins (*on*-switching brightness increased by +13 %), while the V151L mutants switch *on* more efficiently (+50%), mostly originating from a higher *on*-switching quantum yields in the V151L variants. The reduced *on*-to-*off* switching quantum yield in the V151L variants might be attributed to the presence of the bulky leucine raising the excited state energy barrier between the *on*- and *off*-states.

*Supplementary text S4: Modulation of conformational off-state heterogeneity*

In addition to the well documented *trans1* conformation of the *off*-state chromophore [1, 19], a second conformation, *trans2*, has been reported for crystalline rsEGFP2 [3, 30]. Here, we confirm the observation of *trans2* in rsEGFP2 (Figure 1) and provide evidence for its existence in rsFolder2 (Supplementary figure S6). We show that the occupancy of *trans1* and *trans2* conformations can be controlled by modifying the steric constraints imposed by the side-chain of residue 151, a residue retracting transiently during *off*-to-*on* photoswitching [1]. Upon *on*-to-*off* switching, only *trans1* forms in the V151A rsEGFP2 variant, whereas only *trans2* seems to be occupied in the V151L variant (Figure 2).

Interestingly, only the *trans2* conformation is observed in rsFolder, although it has a valine at position 151 [19]. rsFolder displays a phenylalanine at position 146 instead of a tyrosine in rsEGFP2 and rsFolder2 that cannot form an H-bond to the critical His149 in the *off*-state [3, 19]. Consequently, it seems plausible that His149 is free to form an H-bond with the protonated *off*-state chromophore that during on-to-*off* switching then adopts the *trans2* conformer, instead of the *trans1* conformer that is not H-bonded to His149 in rsEGFP2 and rsFolder2. On the other hand, rsGreen0.7 [31] features a phenylalanine at position 146, like rsFolder, but an alanine at position 151, and its *off*-state shows essentially the *trans1* conformer. Thus, access to *trans1* or *trans2* is controlled by residues located either on the *cis* or the *trans* sides of the chromophore pocket, although it appears that the presence of an alanine at position 151 has a dominant effect on the presence of the *trans1* isomer.

***Supplementary tables and figures***

**Table S1 :** Cryo-crystallographic synchrotron data collection and refinement statistics of rsEGFP2-V151A, rsEGFP2-V151L and rsFolder2 structures in their *on*- and *off*-states solved from

| Data collection and processing | | | | | | |
|---|---|---|---|---|---|---|
| Dataset | rsEGFP2 V151A *on* | rsEGFP2 V151A *off* | rsEGFP2 V151L *on* | rsEGFP2 V151L *off* | rsFolder2 *on* | rsFolder2 *off* |
| PDB entry | 7O7D | 7O7C | 7O7E | 7O7H | 7AMB | 7AMF |
| Illumination (488 nm) | No | Yes | No | Yes | No | Yes |
| Space group | $P2_12_12_1$ | $P2_12_12_1$ | $P2_12_12_1$ | $P2_12_12_1$ | C2 | C2 |
| Unit cell parameters | | | | | | |
| a (Å) | 60.0 | 51.2 | 51.1 | 51.3 | 142.3 | 142.3 |
| b (Å) | 62.1 | 60.3 | 62.5 | 61.2 | 134.5 | 134.9 |
| c (Å) | 69.4 | 66.6 | 70.4 | 70.3 | 51.6 | 51.7 |
| β (°) | | | | | 106.1 | 106.0 |
| Collected frames | 1050 | 547 | 794 | 950 | 3700 | 3700 |
| Observations | 166,004 (12,182)* | 89,498 (6,540) | 91,956 (14,602) | 84,740 (13,701) | 389,403 (15,763) | 408,905 (19,013) |
| Resolution (Å) | 46.28 - 1.4 (1.44-1.4) | 40.57 - 1.55 (1.59-1.55) | 41.38 - 1.8 (1.91-1.8) | 46.16 - 1.7 (1.81-1.7) | 47.94 – 1.63 (1.66-1.63) | 48.07 -1.63 (1.66-1.63) |
| $R_{meas}$ (%) | 4.8 (42.5) | 4.5 (60.6) | 6.5 (46.6) | 4.7 (42.8) | 6.5 (73.0) | 7.2 (80.7) |
| $CC_{½}$ (%) | 99.9 (85.3) | 99.9 (70.7) | 99.9 (91.6) | 99.9 (87.1) | 99.9 (74.7) | 99.8 (67.9) |
| I/σI | 17.26 (3.23) | 14.82 (2.26) | 14.17 (2.72) | 15.5 (2.9) | 13.2 (1.8) | 10.3 (1.5) |
| Completeness (%) | 97.4 (96.7) | 97.0 (98.4) | 98.3 (98.3) | 97.2 (98.5) | 99.7 (99.7) | 99.8 (99.5) |
| Multiplicity | 3.9 (3.9) | 3.0 (3.0) | 4.3 (4.3) | 3.5 (3.5) | 3.5 (3.1) | 3.5 (3.3) |
| Refinement statistics | | | | | | |
| Resolution (Å) | 46.28 - 1.4 (1.43-1.40) | 40.57 - 1.55 (1.60-1.55) | 41.38 - 1.8 (1.88-1.80) | 46.16 - 1.7 (1.77-1.70) | 47.98 – 1.63 | 48.12- 1.63 |
| Number of reflections | 42,980 (2,669) | 29,619 (2,528) | 21,205 (2,482) | 24,184 (2,539) | 115,478 (2000) | 116,053 (2000) |
| $R_{free}$ (%) | 17.90 (25.39) | 19.61 (28.60) | 20.76 (29.05) | 19.70 (26.05) | 19.0 | 18.4 |
| $R_{work}$ (%) | 15.21 (21.80) | 16.40 (24.46) | 15.92 (24.97) | 15.70 (22.17) | 15.1 | 15.5 |
| Number of protein atoms | 2,066 | 2,096 | 1,977 | 2,129 | 8,108 | 8,190 |
| Number of solvent atoms | 353 | 275 | 317 | 298 | 1014 | 949 |
| B-factor protein ($Å^2$) | 16.11 | 19.12 | 29.12 | 29.66 | 24.97 | 25.44 |
| r.m.s.d bond lengths (Å) | 0.006 | 0.008 | 0.006 | 0.01 | 0.011 | 0.012 |
| r.m.s.d angles (°) | 0.902 | 0.997 | 0.847 | 1.031 | 1.764 | 1.996 |
| Ramachandran favored (%) | 99.12 | 98.68 | 98.70 | 97.84 | 98.22 | 98.35 |
| Ramachandran allowed (%) | 0.88 | 1.32 | 1.30 | 2.16 | 1.67 | 1.54 |
| Ramachandran outliers (%) | 0.00 | 0.00 | 0.00 | 0.00 | 0.11 | 0.11 |
| Rotamer outliers (%) | 1.29 | 1.28 | 1.79 | 2.08 | 2.10 | 2.84 |
| C-beta outliers | 0 | 0 | 0 | 0 | 2 | 0 |
| Clashscore | 4.32 | 5.66 | 3.29 | 5.81 | 8.24 | 9.24 |

* Values in brackets correspond to the highest resolution shell

**Table S2 :** RT SFX data collection and refinement statistics of the structures of rsEGFP2 and its V151A and V151L variants in their *off*-states

| Dataset | Parental rsEGFP2 *off* | rsEGFP2-V151A *on* | rsEGFP2-V151A *off* | rsEGFP2-V151L *off* |
|---|---|---|---|---|
| PDB entry | 7O7U | 7O7V | 7O7X | 7O7W |
| Data collection and processing | | | | |
| Space group | $P2_12_12_1$ | $P2_12_12_1$ | $P2_12_12_1$ | $P2_12_12_1$ |
| Unit cell parameters<br>a (Å)<br>b (Å)<br>c (Å) | <br>51.7± 0.1<br>63.1 ± 0.2<br>73.5 ± 0.2 | <br>51.9± 0.2<br>62.7 ± 0.3<br>72.0 ± 0.4 | <br>51.8 ± 0.2<br>62.7 ± 0.2<br>71.6 ± 0.3 | <br>51.8 ± 0.2<br>62.9 ± 0.2<br>71.9 ± 0.3 |
| Collected frames | | | | |
| Hits | | | | |
| Indexed images | 80,605 | 17,538 | 10,794 | 4,930 |
| Resolution (Å) | 17– 1.70 (1.74 – 1.70) | 24 – 1.9 (1.95 – 1.90) | 24– 1.95 (2.00 – 1.95) | 24– 2.10 (2.15 –2.10) |
| Observations | 22,894,347 (991,284) | 2,916,477 (100,602) | 2,044,776 (75,714) | 778,314 (28,668) |
| Unique reflections | 27,394 (1,810) | 19,739 (1,293) | 18,181 (1,176) | 14,734 (958) |
| $R_{split}^{\#}$ (%) | 6.8 (56.3) | 6.3 (73.6) | 7.7 (58.4) | 14.0 (53.4) |
| CC* | 0.999 (0.924) | 0.999 (0.891) | 0.999 (0.904) | 0.996 (0.891) |
| I / σ(I) | 10.0 (1.9) | 7.8 (1.4) | 7.0 (1.8) | 4.9 (1.9) |
| Completeness (%) | 100 (100) | 100 (100) | 100 (100) | 100 (100) |
| Multiplicity | 835 (548) | 148 (78) | 112 (64) | 53 (30) |
| Refinement statistics | | | | |
| Refinement strategy | Classical refinement | Classical refinement | Classical refinement | Classical refinement |
| Resolution (Å) | 17– 1.70 (1.74 – 1.70) | 24 – 1.9 (1.95 – 1.90) | 31 – 1.95 (2.00 – 1.95) | 31– 2.10 (2.15 –2.10) |
| Number of reflections | 26368 (2,661) | 19,091 (2,543) | 17,600 (2,739) | 14,284 (2,661) |
| $R_{free}$ | 21.5 (38.0) | 19.2 (27.14) | 20.1 (31.0) | 21.3 (22.0) |
| $R_{work}$ | 18.0 (37.0) | 14.9 (21.16) | 15.7 (26.0) | 17.3 (29.0) |
| Number of protein atoms | 2,443 | 2005 | 2,182 | 2,193 |
| Number of ligand atoms | 60 | 20 | 40 | 40 |
| Number of water atoms | 177 | 111 | 125 | 58 |
| B-factor protein (Å$^2$) | 22.81 | 48.5 | 43.41 | 38.97 |
| r.m.s.d. bond lengths (Å) | 0.01 | 0.01 | 0.008 | 0.008 |
| r.m.s.d. angles (°) | 1.728 | 1.609 | 1.547 | 1.566 |
| Ramachandran favoured | 97.00 | 97.90 | 97.83 | 97.41 |
| Ramachandran allowed | 3.00 | 2.10 | 2.17 | 2.59 |
| Ramachadran outliers | 0.00 | 0.00 | 0.00 | 0.00 |
| Rotamer outliers | 4.01 | 1.80 | 1.22 | 0.81 |
| C-beta outliers | 0 | 0 | 0 | 0 |
| Clashscore | 6.23 | 2.51 | 5 | 6.78 |

**Table S3** : Chromophore dihedral angles from crystallographic models and after computational geometry optimization

| Method | Protein | State | PDB ID | Conformer | * φ (°) | # τ (°) |
|---|---|---|---|---|---|---|
| Synchrotron cryo-crystallography | rsEGFP2-V151A | *off*-state | 7O7C | *cis* | 174 | -166 |
| | | | 7O7C | *trans1* | -54 | 16 |
| | | *on*-state | 7O7D | *cis* | 174 | -166 |
| | rsEGFP2-V151L | *off*-state | 7O7H | *cis* | 171 | -169 |
| | | | 7O7H | *trans2* | 7 | -3 |
| | | *on*-state | 7O7E | *cis* | 171 | -169 |
| | rsFolder2 | *on*-state | 7AMB | *cis* | 175 | -178 |
| | | *off*-state | 7AMF | *cis* | 170 | -176 |
| | | | 7AMF | *trans1* | -44 | 17 |
| | | | 7AMF | *trans2* | 7 | -4 |
| | Cl-rsEGFP2 [30] | *on*-state | 6PFR | expanded *cis* | 167 | -166 |
| | | | 6PFS | contracted *cis* | 142 | -169 |
| | | *off*-state | 6PFT | expanded *trans* | -32$ | 2 |
| | | | 6PFU | contracted *trans* | 43 | -30 |
| RT SFX | rsEGFP2 | *off*-state | 7O7U | *cis* | 168 | -167 |
| | | | 7O7U | *trans1* | -48 | 7 |
| | | | 7O7U | *trans2* | 7 | -4 |
| | rsEGFP2-V151A | *off*-state | 7O7X | *cis* | 173 | -172 |
| | | | 7O7X | *trans1* | -50 | 9 |
| | | *on*-state | 7O7V | *cis* | 178 | -170 |
| | rsEGFP2-V151L | *off*-state | 7O7W | *cis* | 173 | -151 |
| | | | 7O7W | *trans2* | 10 | -18 |
| Comput. B3LYP-D3 geometry optimization | rsEGFP2 | *off*-state | (6T39) & | *trans1* | -31 | -5 |
| | | *off*-state | (6T39) & | *trans2* | -7 | -10 |
| | rsEGFP2-V151A | *off*-state | (6T39) & | *trans1* | -27 | -6 |
| | rsEGFP2-V151L | *off*-state | (6T39) & | *trans2* | -8 | -10 |

*CA2-CB2-CG2-CD2 (for numbering see Supplementary figure 16c in [1])
#C2-CA2-CB2-CG2
$CA2-CB2-CG2-CD1 (due to the flip of the chromophore around CB2-CG2 in PDB ID 6PFT)

&starting coordinates of the computational models were taken from PDB 6T39

**Table S4.** Results of XMCQDPT2 excited-state calculations comparing *trans1* and *trans2*. The calculations employed the SA3-CASSCF(2e,2MO) and SA4-CASSCF(4e,3MO) (indicated as CT[a)]) zero-order approximation.

| Chromophore conformation | Geometry | Excitation energy (eV) and transition dipole moment (Hartree*Bohr, indicated in brackets) of $S_0$-Sn | | |
|---|---|---|---|---|
| | | $S_0$-$S_1$ | $S_0$-$S_2$ (CT Y149) | $S_0$-$S_3$ |
| *Trans1* | V151, $S_0$-min | 2.98 (2.44) | - | 5.07 (0.60) |
| | V151, R(O-H)=1.5 Å | 2.59 (2.61) | - | 4.82 (0.50) |
| | A151, $S_0$-min | 2.98 (2.50) | - | 5.06 (0.65) |
| | V151, $S_0$-min, CT[a)] | 2.82 (2.67) | 3.92(0.05) | 5.00 (0.69) |
| *Trans2* | V151, $S_0$-min | 2.90 (2.77) | - | 4.95 (2.76) |
| | V151, R(O-H)=1.5 Å | 2.55 (3.03) | - | 4.71 (0.61) |
| | L151, $S_0$-min | 2.86 (2.78) | - | 4.94 (0.75) |
| | V151, $S_0$-min, CT[a)] | 2.78 (3.00) | 4.74 (0.002) | 5.04 (0.79) |

a) The SA4-CASSCF(4e,3MO) calculations.

**Table S5.** Vibrational energy levels ($cm^{-1}$) relative to the $S_0$-min energy and energies of the vibronic transition computed for one-dimensional electronic potentials presented in Supplementary figure S9.

| Chromophore conformation | $S_0$ state | | $S_1$ state | | $S_0$-$S_1$ vibronic transitions | |
|---|---|---|---|---|---|---|
| *Trans1* | $S_0$-min | 0 | $S_1$-min | 24071 | VEE[a)] | 24071 (2.98 eV) |
| | *v*=0 | 1724 | *v*=0 | 24390 | 0-0 | 22666 ( 2.81 eV) |
| | *v*=1 | 4333 | *v*=1 | 25655 | 0-1 | 23931 (2.97 eV) |
| | *v*=2 | 6146 | *v*=2 | 27211 | 0-2 | 25487 (3.04 eV) |
| | *v*=3 | 7952 | *v*=3 | 28936 | 0-3 | 27212 |
| | *v*=4 | 9867 | *v*=4 | 30833 | 0-4 | 29109 |
| | *v*=5 | | *v*=5 | 32902 | 0-5 | 31178 |
| *Trans2* | $S_0$-min | 0 | $S_1$-min | 23445 | VEE [a)] | 23445 (2.91 eV) |
| | *v*=0 | 965 | *v*=0 | 22027 | 0-0 | 21062 (2.61 eV) |
| | *v*=1 | 1541 | *v*=1 | 23194 | 0-1 | 22229 (2.76 eV) |
| | *v*=2 | 2472 | *v*=2 | 24196 | 0-2 | 23231 (2.88 eV) |
| | *v*=3 | 3575 | *v*=3 | 25237 | 0-3 | 24272 |
| | *v*=4 | 4836 | *v*=4 | 26476 | 0-4 | 25511 |
| | *v*=5 | | *v*=5 | | | |

a) VEE stands for vertical excitation energy

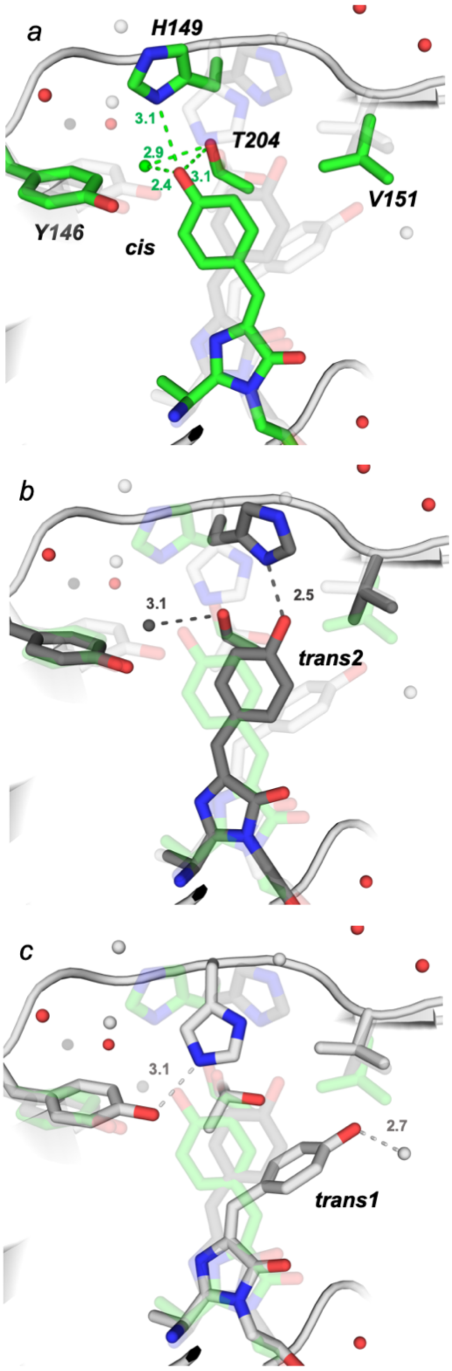

**Supplementary figure S1**: Environment of the chromophore in its *cis* (a; green) , *trans2* (b; dark grey) and *trans1* (c; light grey) conformation in the parental rsEGFP2 *off*-state structure (PDB entry 7O7U). Water molecules belonging to all three chromophore conformations are shown in red. Hydrogen-bond lengths are in Ångstrom.

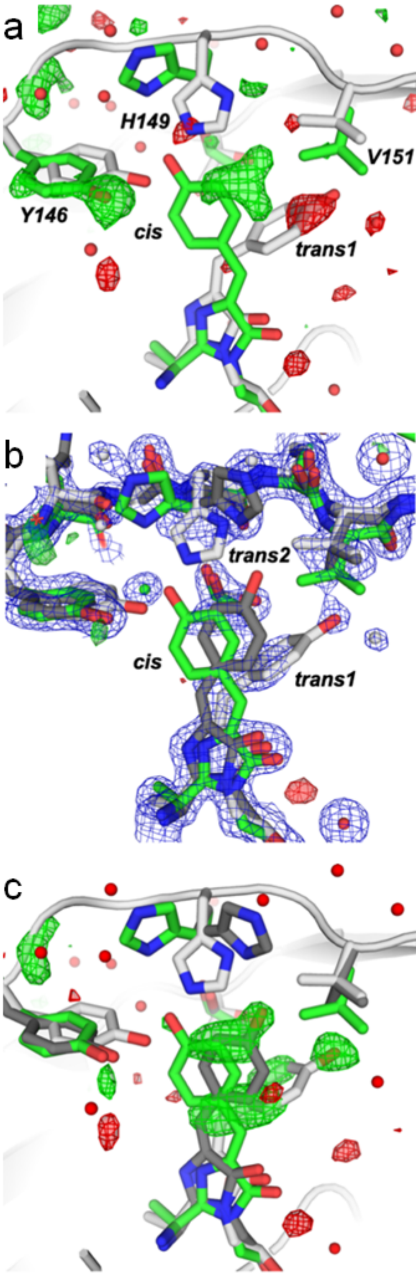

**Supplementary figure S2**: Structure of parental rsEGFP2 in its *off*-state solved from RT SFX data. (a) Residual $F_{obs} – F_{calc}$ map (green, 3σ ; red, -3σ) of the parental rsEGFP2 in the *off*-state calculated with a model containing a *cis* (light green) and a *trans1* chromopore (light grey) at an occupancy of 30 and 70%, respectively. (b) Final model of parental rsEGFP2 in the *off*-state (PDB entry 7O7U) with occupancies of *cis* (green), *trans1* (light grey) and *trans2* (dark grey) chromophore conformations of 30%, 40% and 30%, respectively. Superimposed are the 2 $F_{obs} – F_{calc}$ (blue, 1 σ) and $F_{obs} – F_{calc}$ (green, 3 σ ; red, -3 σ) maps. (c) $F_{obs} – F_{calc}$ omit map (hydroxybenzylidene moiety omitted from the model; green, 3 σ ; red, -3 σ), overlaid with the model of parental rsEGFP2 in the *off*-state (PDB entry 7O7U).

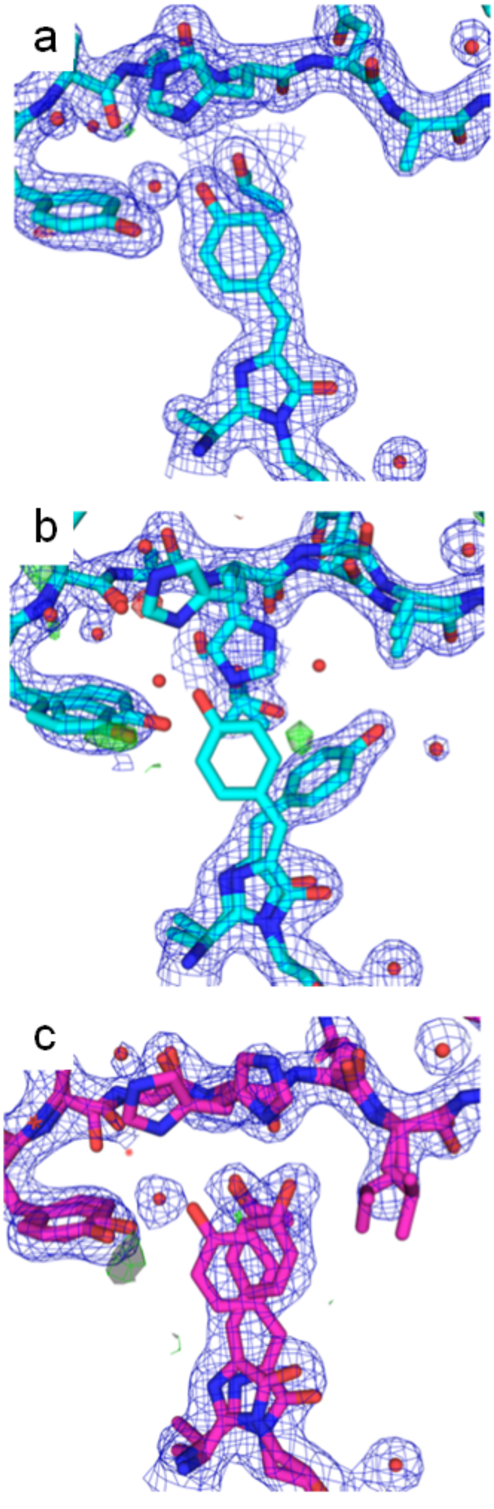

-

**Supplementary figure S3:** Zoom on the chromophore (p-hydroxybenzylidene imidazolinone, p-HBI) region in crystal structures of (a) the rsEGFP2-V151A variant in its *on-* (PDB entry 7O7V) and (b) *off*-state (PDB entry 7O7X) and (c) the V151L variant in its *off*-state (PDB entry 7O7W) solved from SFX data collected at room temperature. $2F_{obs} - F_{calc}$ ($1\sigma$) and $F_{obs} - F_{calc}$ (+/- $3\sigma$) electron density maps are shown in blue and green/red, respectively.

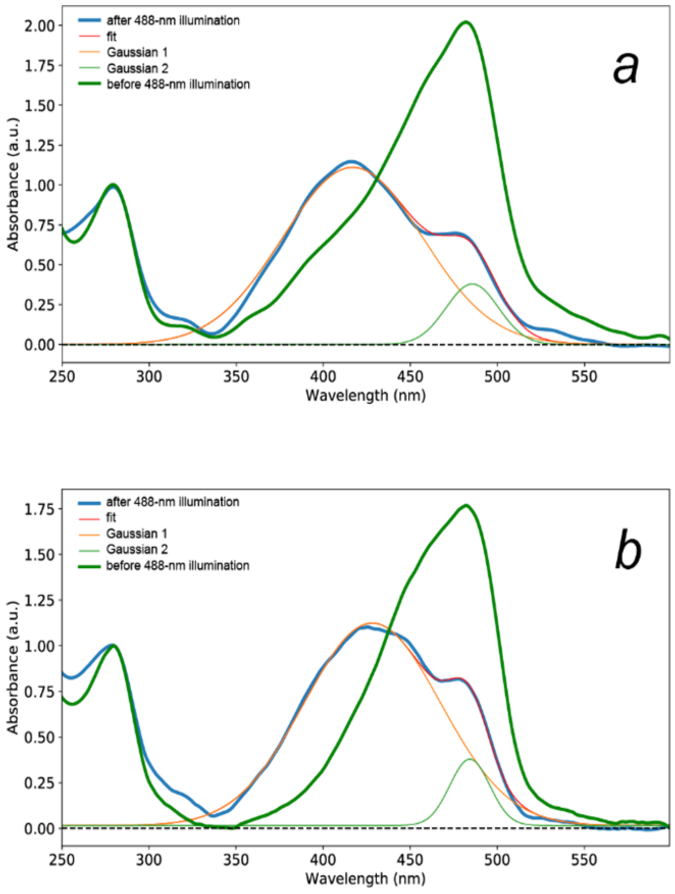

**Supplementary figure S4**: Absorption spectra of rsEGFP2-V151A (*a*) and -V151L (*b*) microcrystals before (green) and after having been pre-illuminated (blue) at a laser power of nominally 200 mW within a custom-made device [9] during an SFX experiment at SACLA (July 2018) after background subtraction, smoothening with a Savitzky-Golay filter and normalization at 280 nm. The spectrum after 488 nm illumination was modeled by a sum of two Gaussians (Gaussian 1 at 400 nm showed in orange and Gaussian 2 at 482 nm showed in light green). The absorbance of Gaussian 2 at 482 nm relative to the absorbance at 482 nm of the spectrum before 488 nm illumination (assumed to correspond to 100% *on*-state) indicates that 85% (77%) were switched to the *off*-state for rsEGFP2-V151A (V151L) and 15% (23%) remained in the *on*-state.

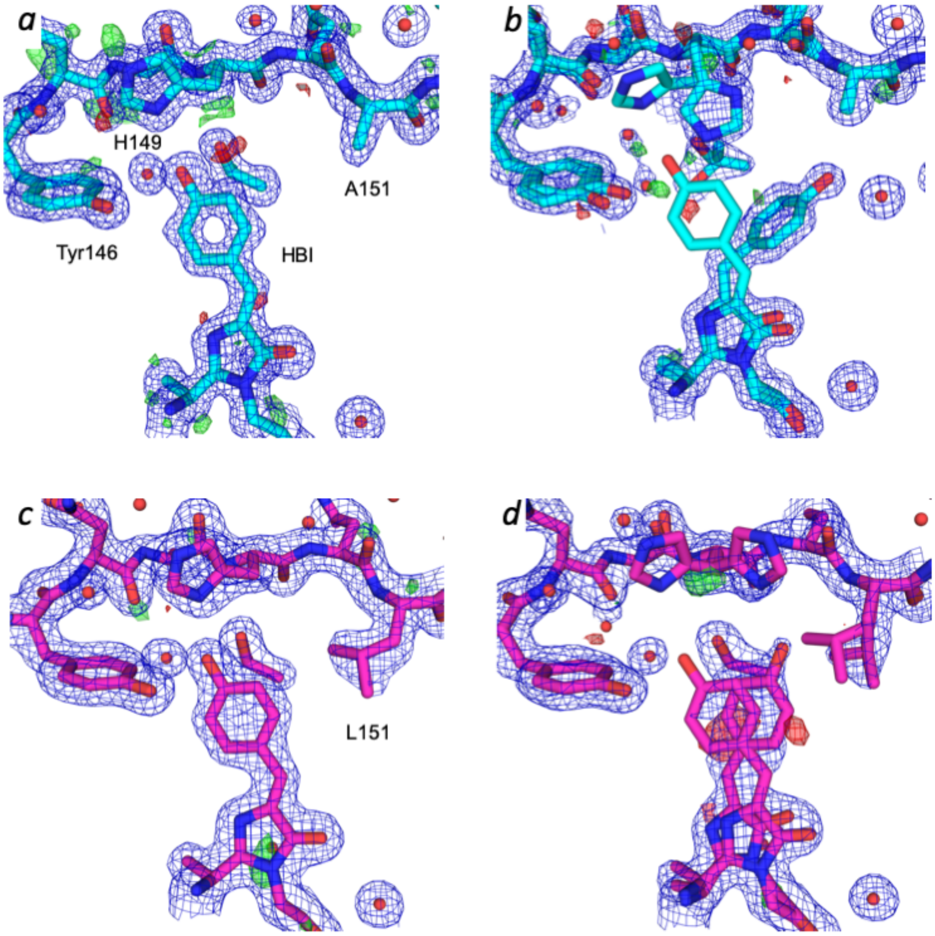

**Supplementary figure S5**: Zoom on the chromophore (p-hydroxybenzylidene imidazolinone, p-HBI) region in crystal structures of rsEGFP2 variants V151A (cyan) and V151L (purple) in their *on*- and *off*-states solved from synchrotron data collected at cryo-temperature. (*a, b*) Electron density maps (mesh) and models of the rsEGFP2-V151A variant in the (*a*) *on*- (PDB entry 7O7D) and (*b*) *off*-state (PDB entry 7O7C). (*c, d*) Electron density maps (mesh) and models of the rsEGFP2-V151L variant in the (*c*) *on*- (PDB entry 7O7E) and (*d*) *off*-state (PDB entry 7O7H). In (*d*) an alternate conformation of the chromophore corresponding to the *cis* isomer of the *on*-state remains in the *off*-state. $2F_{obs}$-$F_{calc}$ (1 σ) and $F_{obs}$-$F_{calc}$ (+/- 3 σ) electron density maps are shown in blue and green/red, respectively.

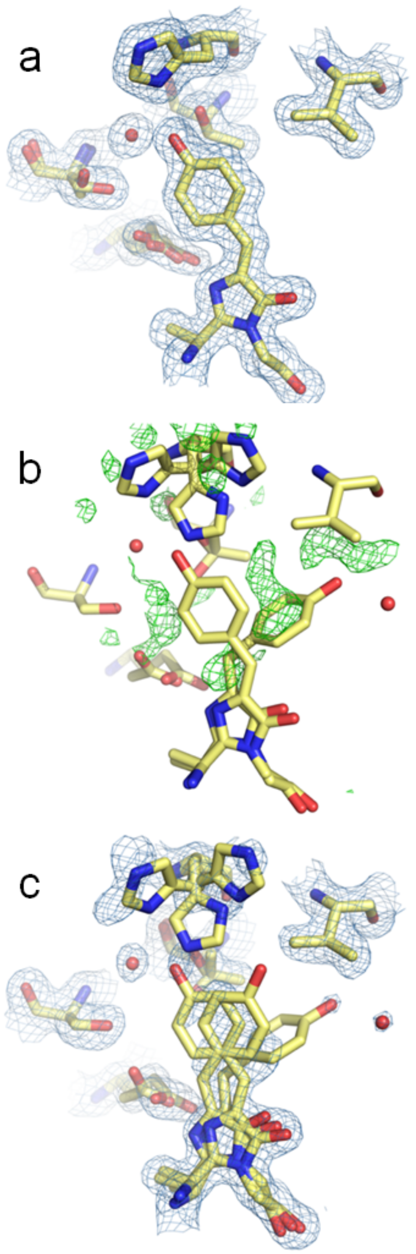

**Supplementary figure S6:** rsFolder2 *on*- and *off*-state structures featuring evidence for a low occupancy *trans2* conformation in monomer A, derived from synchrotron cryo-crystallographic data. Electron density $2F_{obs}$-$F_{calc}$ maps are shown as a blue mesh at 1σ and the $F_{obs}$-$F_{calc}$ difference map is shown as a green mesh at 3 σ. (a) Active site environment for the *on*-state structure (PDB entry 7AMB), with a fully occupied *cis* chromophore. (b) active site for the *off*-structure, where only *cis* and *trans1* conformers have been included in the model for refinement. NCS averaged difference map is shown in green, indicating the presence of the *trans2* conformer. (c) refined *off*-state model (PDB entry 7AMF) for monomer A, with *cis* (60%), *trans1* (30%) and *trans2* (10%).

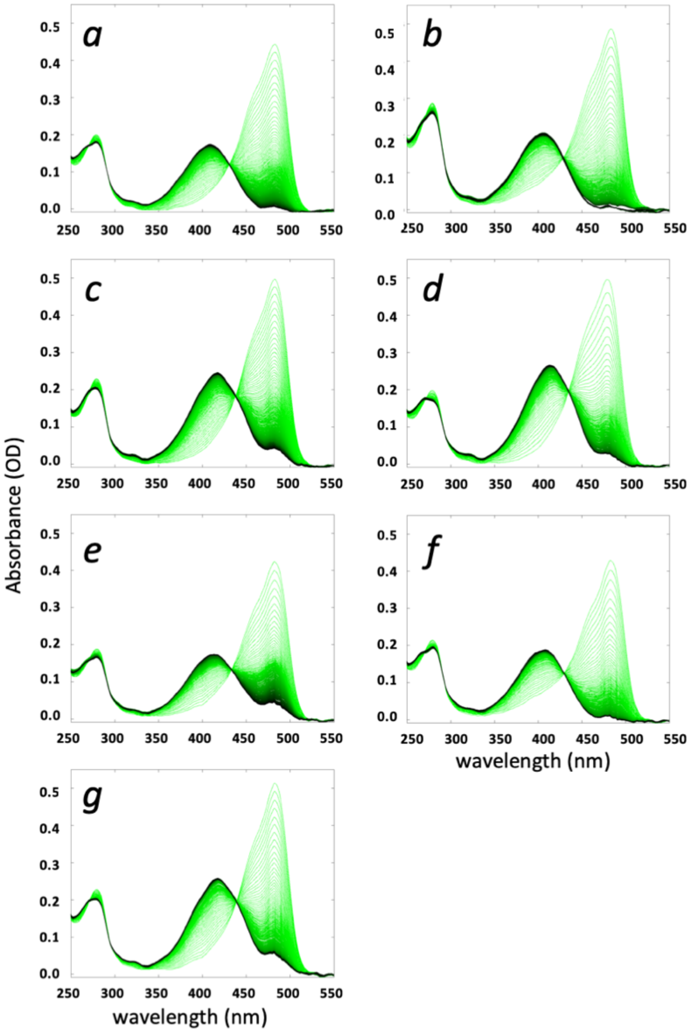

**Supplementary figure S7**: Absorption spectra along *on*-to-*off* photoswitching of the RSFPs presented in this study measured in solution in HEPES buffer at pH 7.5. Photoswitching series of spectra are color-coded from light green (*on*) to black (*off*) for rsEGFP2 (a), rsEGFP2-V151A (b), rsEGFP2-V151L (c), rsFolder (d), rsFolder2 (e), rsFolder2-V151A (f) and rsFolder2-V151L (g)

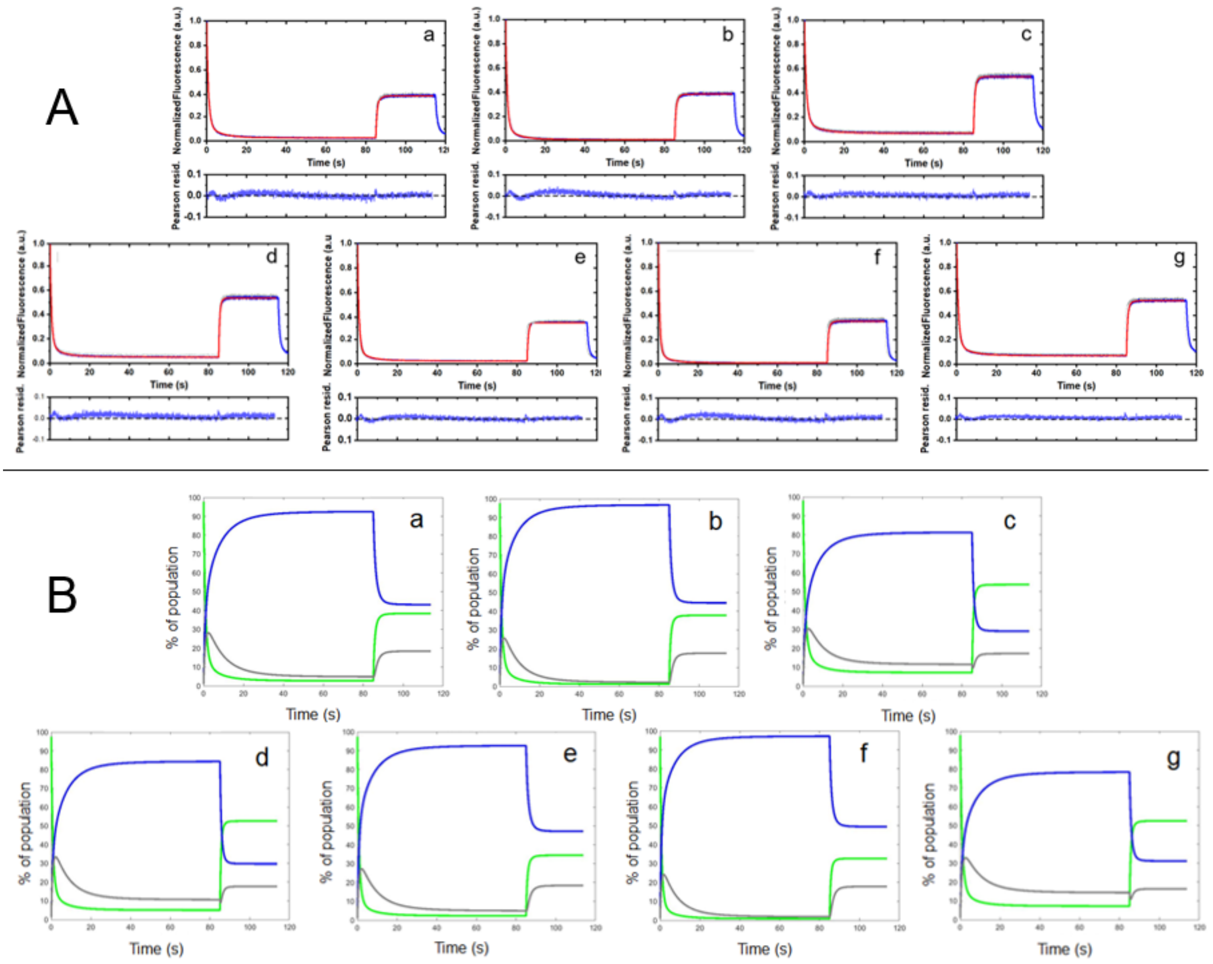

**Supplementary figure S8:** Switching curves (A) and concentration plots (B) for rsEGFP2 (a), rsEGFP2-V151A (b), rsEGFP2-V151L (c), rsFolder (d), rsFolder2 (e), rsFolder2-V151A (f) and rsFolder2-V151L (g).

A: Fluorescence switching curves. The mean of six measurements is represented in blue with a standard deviation in grey. Data was fitted with a model (red) that includes an additional dark state. Pearson residuals below each graph demonstrate the goodness of fit.

B: Concentration plots. The percentage of protein population are shown in the *on*-state (green), the *off*-state (blue) and in the additional dark state (grey) along the switching cycle. The fluorescence profile of the full switching cycle was fitted with a kinetic model including two reversible dark-states.

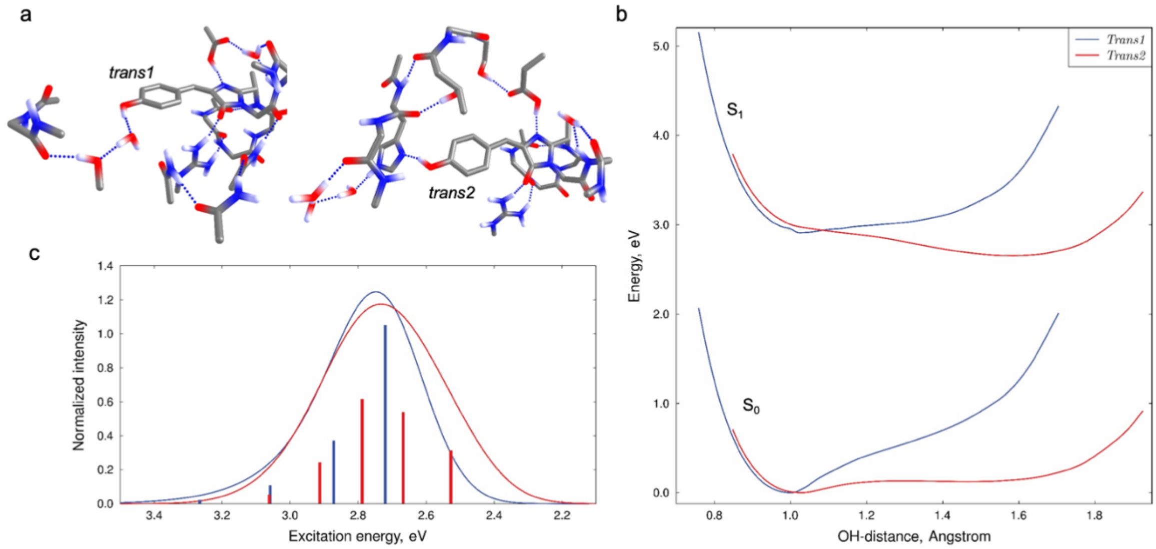

**Supplementary figure S9**. The phenolic group OH-stretching potentials in the ground and excited states and corresponding vibronic band of the $S_0$-$S_1$ transition. (a) demonstrates models of the chromophore employed in computations. (b) presents the $S_0$ and $S_1$ energies computed with the XMCQDPT2-SA3-CASSCF(2e,2MO) method at the geometries of the relaxed OH-stretching energy scans computed in the ground state with the B3LYP-D3/cc-pvdz method. The resulting $S_0$ and $S_1$ OH stretching potentials were used to simulate the vibronic band shown in (c). The energies and probability density for vibronic states for a one-dimensional potential were computed using a numerical procedure. The energies are collected in Supplementary table S5. Broadening of the stick spectra was done with Gaussian functions (1000 $cm^{-1}$ FWHM); the intensities were normalized to the $S_0$-$S_1$ transition dipole moments computed at the $S_0$ equilibrium geometries.

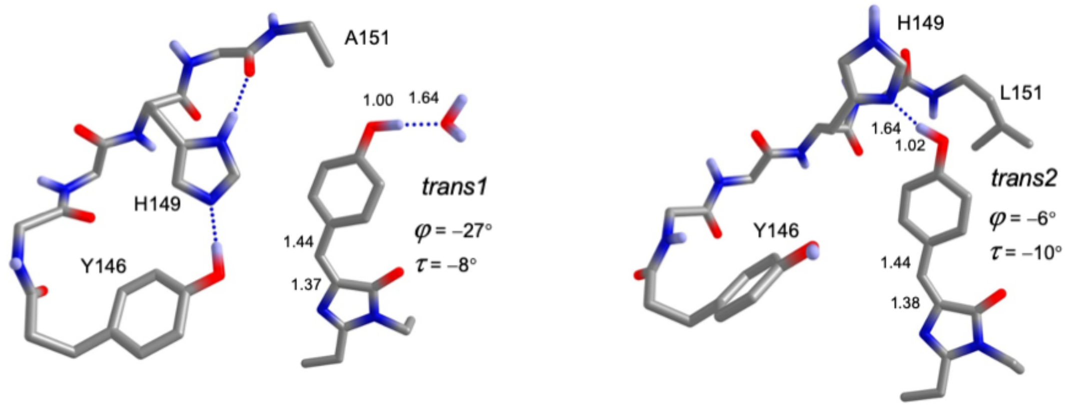

**Supplementary figure S10:** Optimized geometries obtained for the active sites containing the V151A (left) and V151L (right) substitutions. The initial coordinates of the atoms were obtained by substituting the Val151 side chain of parental rsEGFP2 (PDB entry 6T39[3]) by Ala or Leu side chains. The geometry was optimized with the B3LYP-D3/cc-pvdz method. At the optimized geometries, the excitation energies were computed with the XMCQDPT2/cc-pvdz method (SA3-CASSCF(2,2) wave function). The excitation energies and transition moments are compared with those obtained for rsEGFP2 active-site models are presented in Supplementary table S4.

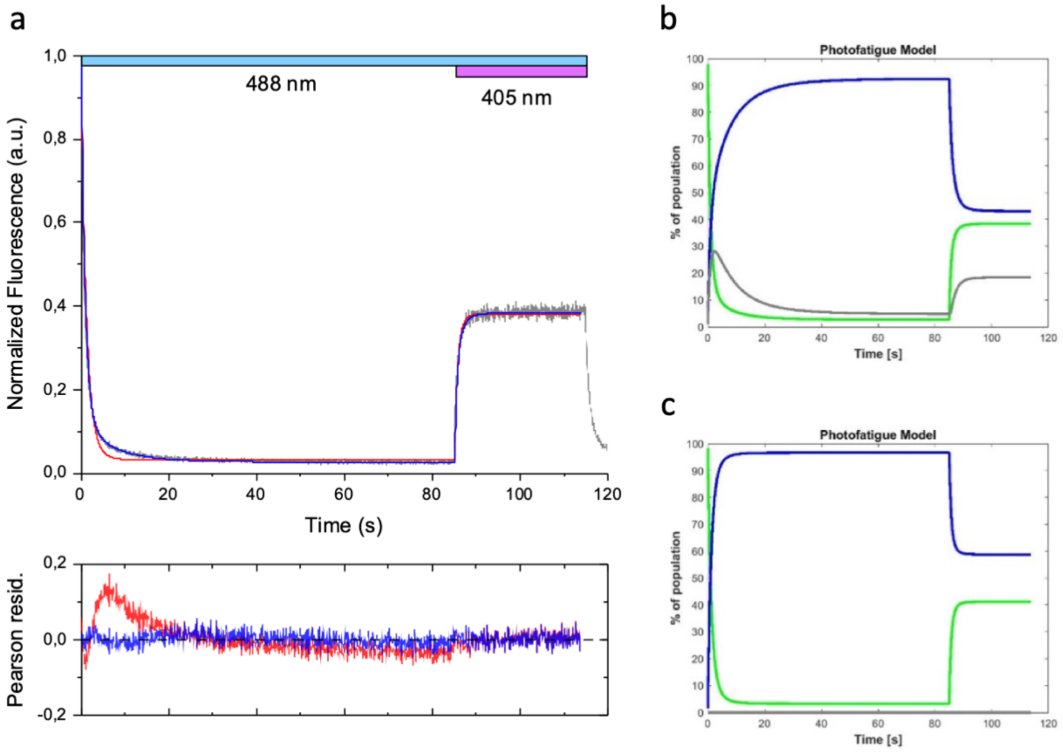

**Supplementary figure S11**: Additional dark state during photoswitching of parent rsEGFP2. (a) Fitted switching profiles using a model that includes (blue) or does not include (red) an additional dark state, overlaid on experimental switching profile (grey); Pearson residuals for each fit (bottom panel). (b, c) Fitted populations along time for both models (*on*-state in green, *off*-state in blue and the additional dark state in grey).

.

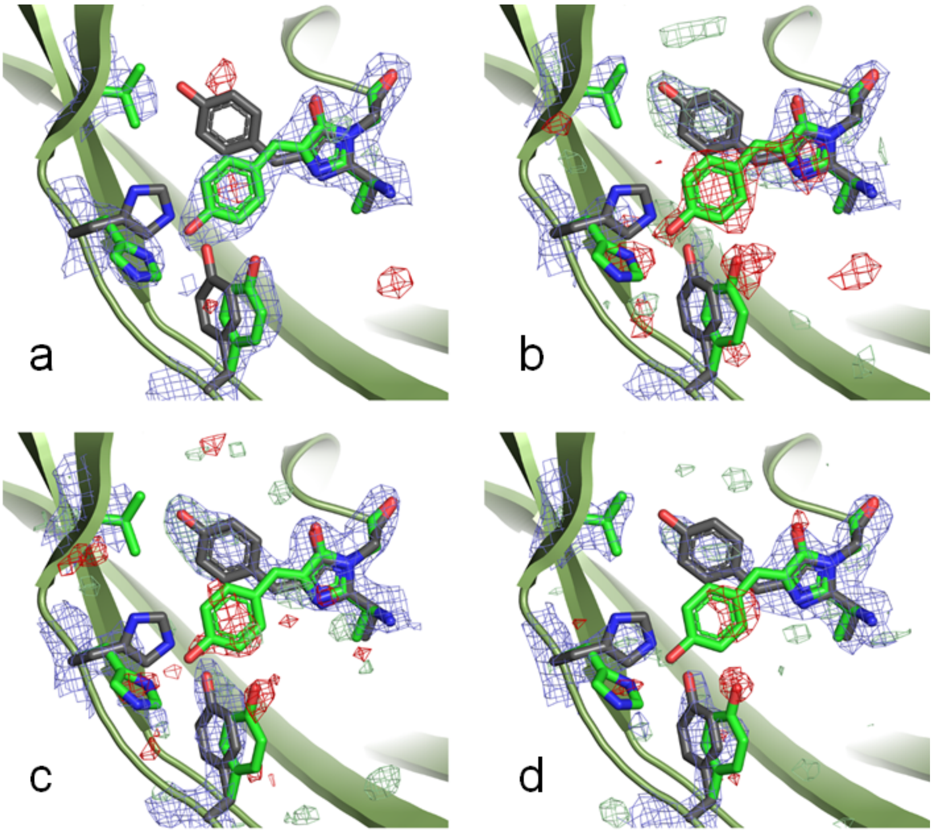

**Supplementary figure S12:** Structures and electron density maps of rsEGFP2 photoswitched at different power densities. Each crystal was photoswitched during ~30 s at 100 mW/cm$^2$ (a), 200 mW/cm$^2$ (b), 400 mW/cm$^2$ (c) or 800 mW/cm$^2$ (d) and immediately cryoprotected and flash-cooled. $2F_{obs} - F_{calc}$ electron density maps (blue) are shown at 1.5 σ, $F_{obs} - F_{calc}$ difference electron density maps are shown at 3 σ (green) and -3σ (red).

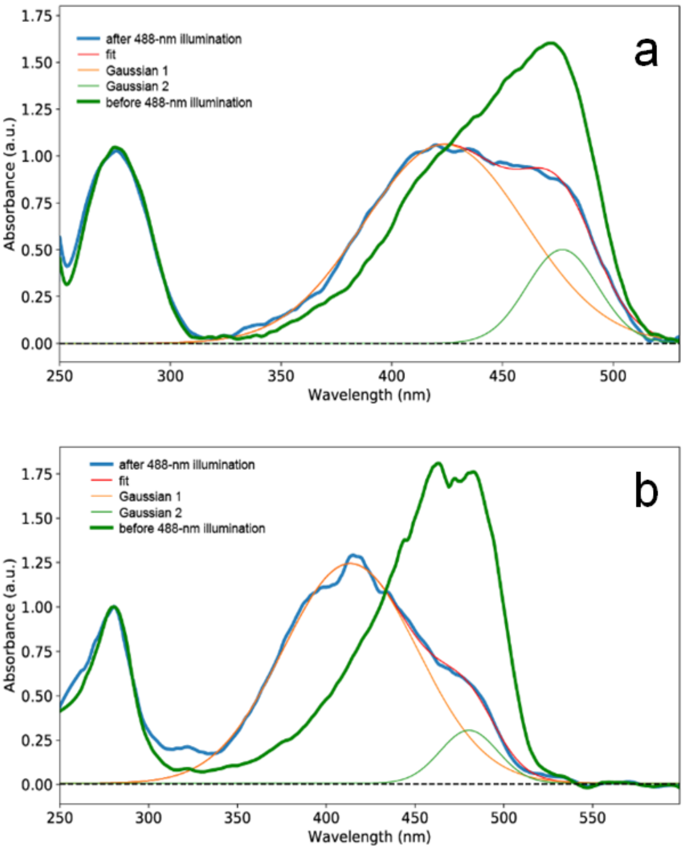

**Supplementary figure S13**: Absorption spectra of parental rsEGFP2 microcrystals before (green) and after having been pre-illuminated (blue) at a laser power of nominally 200 mW within a custom-made device [9] during SFX experiments at the LCLS in June 2016 (LM47) (a) and at SACLA in July 2015 [3] (b) after background subtraction, smoothening with a Savitzky-Golay filter and normalization at 280 nm. The spectrum after 488 nm illumination was modeled by a sum of two Gaussians (Gaussian 1 at 400 nm and Gaussian 2 at 482 nm). The absorbance at 482 nm as fitted by Gaussian 2 relative to the absorbance at 482 nm of the spectrum before 488 nm illumination (assumed to correspond to 100% *on*-state) indicates that 66% (84%) were switched to the *off*-state in panel a (b) and 34% (16%) remained in the *on*-state.

[17] D. de Sanctis, A. Beteva, H. Caserotto, F. Dobias, J. Gabadinho, T. Giraud, A. Gobbo, M. Guijarro, M. Lentini, B. Lavault, T. Mairs, S. McSweeney, S. Petitdemange, V. Rey-

Bakaikoa, J. Surr, P. Theveneau, G. A. Leonard, C. Mueller-Dieckmann, *Journal of synchrotron radiation* **2012**, *19*, 455-461.
[18] W. Kabsch, *Acta crystallographica. Section D, Biological crystallography* **2010**, *66*, 133-144.
[19] M. El Khatib, A. Martins, D. Bourgeois, J. P. Colletier, V. Adam, *Scientific reports* **2016**, *6*, 18459.
[20] P. V. Afonine, R. W. Grosse-Kunstleve, N. Echols, J. J. Headd, N. W. Moriarty, M. Mustyakimov, T. C. Terwilliger, A. Urzhumtsev, P. H. Zwart, P. D. Adams, *Acta Crystallographica Section D* **2012**, *68*, 352-367.
[21] W. L. DeLano, *San Carlos, CA, USA: DeLano Scientific* **2002**.
[22] L. Potterton, J. Agirre, C. Ballard, K. Cowtan, E. Dodson, P. R. Evans, H. T. Jenkins, R. Keegan, E. Krissinel, K. Stevenson, A. Lebedev, S. J. McNicholas, R. A. Nicholls, M. Noble, N. S. Pannu, C. Roth, G. Sheldrick, P. Skubak, J. Turkenburg, V. Uski, F. von Delft, D. Waterman, K. Wilson, M. Winn, M. Wojdyr, *Acta Crystallographica Section D* **2018**, *74*, 68-84.
[23] C. J. Williams, J. J. Headd, N. W. Moriarty, M. G. Prisant, L. L. Videau, L. N. Deis, V. Verma, D. A. Keedy, B. J. Hintze, V. B. Chen, S. Jain, S. M. Lewis, W. B. Arendall Iii, J. Snoeyink, P. D. Adams, S. C. Lovell, J. S. Richardson, D. C. Richardson, *Protein Science* **2018**, *27*, 293-315.
[24] M. W. Schmidt, K. K. Baldridge, J. A. Boatz, S. T. Elbert, M. S. Gordon, J. H. Jensen, S. Koseki, N. Matsunaga, K. A. Nguyen, S. Su, T. L. Windus, M. Dupuis, J. A. Montgomery Jr, *Journal of Computational Chemistry* **1993**, *14*, 1347-1363.
[25] aA. D. Becke, *The Journal of Chemical Physics* **1993**, *98*, 5648-5652; bC. Lee, W. Yang, R. G. Parr, *Physical Review B* **1988**, *37*, 785-789; cS. Grimme, J. Antony, S. Ehrlich, H. Krieg, *J Chem Phys* **2010**, *132*, 154104.
[26] A. A. Granovsky, *J Chem Phys* **2011**, *134*, 214113.
[27] R. J. Cave, M. D. Newton, *Chemical Physics Letters* **1996**, *249*, 15-19.
[28] D. E. Woon, T. H. D. Jr., *The Journal of Chemical Physics* **1995**, *103*, 4572-4585.
[29] aE. De Zitter, J. Ridard, D. Thédié, V. Adam, B. Lévy, M. Byrdin, G. Gotthard, L. Van Meervelt, P. Dedecker, I. Demachy, D. Bourgeois, *Journal of the American Chemical Society* **2020**, *142*, 10978-10988; bE. De Zitter, D. Thédié, V. Mönkemöller, S. Hugelier, J. Beaudouin, V. Adam, M. Byrdin, L. Van Meervelt, P. Dedecker, D. Bourgeois, *Nature Methods* **2019**, *16*, 707-710; cR. Berardozzi, V. Adam, A. Martins, D. Bourgeois, *J Am Chem Soc* **2016**, *138*, 558-565.
[30] J. Chang, M. G. Romei, S. G. Boxer, *J Am Chem Soc* **2019**, *141*, 15504-15508.
[31] S. Duwé, E. De Zitter, V. Gielen, B. Moeyaert, W. Vandenberg, T. Grotjohann, K. Clays, S. Jakobs, L. Van Meervelt, P. Dedecker, *ACS nano* **2015**, *9*, 9528-9541.
